# UTR-Diffusion: Conditional Diffusion Modeling for Multi-objective and Constrained UTR Design

**DOI:** 10.64898/2026.08.25.746997

**Authors:** Chuankai Dai, Kengo Sato

## Abstract

**Motivation:** The 5′ untranslated region (UTR) and the start-codon-proximal region of the coding sequence (CDS) jointly influence translation efficiency and local RNA secondary-structure stability, while synonymous codon choices throughout the CDS shape codon adaptation. Because the encoded protein is often predetermined, practical mRNA design must coordinate these quantitative objectives while preserving specified nucleotide sequences and amino-acid identities. Existing generative approaches typically address continuous-valued targeting, explicit sequence constraints, and codon-usage control separately rather than integrating all three within a single model.

**Results:** We present UTR-Diffusion, a diffusion-based framework for 5′ UTR and 5′ UTR–CDS junction design. UTR-Diffusion conditions generation on continuous-valued MRL and MFE targets and supports nucleotide-level constraints, amino-acid-level constraints with synonymous-codon flexibility, and codon-adaptiveness control that modulates the sequence-level codon adaptation index (CAI). Systematic evaluations across dense MRL–MFE target grids showed that generated distributions shifted consistently with both targets, retained substantial diversity, and strictly preserved specified nucleotide sequences and amino-acid identities. Codon-adaptiveness control yielded distinct, monotonically ordered CAI levels that closely followed the specified adaptiveness targets. In comparative benchmarks, UTR-Diffusion outperformed representative existing methods in high-MRL optimization and precise MRL targeting for 5′ UTR design, and achieved higher MRL, less-negative junction MFE, and higher CAI than peptide-preserving baselines in 50-nt 5′ UTR–CDS junction design.

**Availability:** Implementation and trained models for UTR-Diffusion are available at https://github.com/sato-lab-org/utr-diffusion.

**Supplementary information:** Supplementary data are available with the preprint

## Introduction

Messenger RNA (mRNA) plays an important role in therapeutics and synthetic biology, with applications in vaccination, protein replacement, and cell engineering [Qin et al., 2022, Parhiz et al., 2024]. In many applications, the encoded protein is predetermined, so sequence design must tune translation-related and structural properties without altering the amino-acid sequence. For an mRNA encoding a fixed amino-acid sequence, the 5′ UTR and synonymous codon choices throughout the coding sequence (CDS) constitute two major degrees of freedom in sequence design. Throughout the CDS, synonymous codon choices are subject to codon-adaptation objectives; near the translation start site, the same choices additionally interact with the 5′ UTR to shape local RNA folding and start-codon accessibility [Mignone et al., 2002, Tuller et al., 2010, Mauger et al., 2019]. A general computational framework for this design problem should therefore support quantitative target control, preservation of exact nucleotide sequences and amino-acid identities, and graded control of synonymous-codon usage while generating diverse candidate sequences.

Existing in silico approaches address different subsets of these requirements. Predictor-guided search and model-based reinforcement learning can identify biological sequences with high or target-matching scores [Linder et al., 2020, Angermueller et al., 2020]. For 5′ UTR design, Optimus 5-Prime combines a ribosome-loading predictor with sequence search to engineer specified ribosome-loading levels [Sample et al., 2019]. For broader mRNA sequence design, LinearDesign jointly optimizes RNA folding and codon usage, whereas DNAChisel combines user-defined constraints with optimization specifications [Zhang et al., 2023, Zulkower and Rosser, 2020]. These methods are effective for their intended tasks, but rely on task-specific objective and constraint formulations rather than a single learned conditional distribution that supports both extremal optimization and calibrated target tracking under heterogeneous sequence specifications.

Generative models address the need for diverse candidate sets by learning distributions over biological sequences rather than returning isolated optimized solutions. Variational autoencoders and generative adversarial networks have been applied to biological sequence design, and UTRGAN combines GAN-based 5′ UTR generation with optimization toward high translation-related scores [Iwano et al., 2022, Barazandeh et al., 2025]. Large nucleotide language models further demonstrate powerful sequence modeling and structure-aware RNA generation [Zhang et al., 2025]. However, control in these applications is often implemented through unconditional generation, discrete attributes, or downstream reward-based optimization. Consequently, continuous-valued multi-property targeting, exact nucleotide or amino-acid preservation, and graded codon-usage control have largely remained separate capabilities.

Diffusion models provide a promising basis for integrating these capabilities because they combine distributional generation, flexible conditioning, and iterative refinement. RNAdiffusion performs variable-length RNA generation in a latent representation and uses reward guidance to optimize properties including ribosome loading and translation efficiency [Huang et al., 2024], whereas DNA-Diffusion generates 200-bp regulatory DNA sequences under cell-type-specific conditions [DaSilva et al., 2026]. These studies demonstrate the versatility of diffusion-based biological sequence generation, but their published formulations focus primarily on reward-guided optimization or contextual conditioning. A unified framework that directly combines continuous-valued multi-property targeting, exact nucleotide- and amino-acid-level specifications, and graded codon-usage control within 5′ UTR–CDS junction design remains underdeveloped.

To address this gap, we introduce UTR-Diffusion, a sequence-space diffusion framework for 5′ UTR and 5′ UTR–CDS junction design. The model conditions generation on continuous-valued targets for mean ribosome load (MRL), a translation-related measure, and minimum free energy (MFE), a measure of predicted RNA secondary-structure stability. It supports three forms of sequence specification: exact nucleotide-level constraints, amino-acid-level constraints with synonymous-codon flexibility, and codon-adaptiveness control for graded modulation of the sequence-level codon adaptation index (CAI), a measure of synonymous-codon preference [Sharp and Li, 1987]. A RePaint-inspired resampling and re-noising strategy [Lugmayr et al., 2022] enforces nucleotide and amino-acid specifications during reverse diffusion, while an adaptiveness-weighted codon-selection rule controls synonymous-codon preference without retraining the diffusion model. We evaluate the framework across dense MRL–MFE target grids, multiple nucleotide- and amino-acid-level constraint families, and peptide-preserving codon-adaptiveness conditions. We further benchmark UTR-Diffusion in high-MRL optimization and precise MRL targeting for 5′ UTR design, as well as multi-objective MRL–MFE–CAI optimization at the 5′ UTR–CDS junction.

## Methods

### Diffusion Model Framework

The denoising model backbone is implemented with a UNet [Ronneberger et al., 2015] architecture, as illustrated in Figure 1B. Each RNA sequence is encoded as a 4 × *L* matrix whose (*i, j*) entry takes value 1 if nucleotide *i* ∈ {*A, C, G, U*} is present at position *j* and −1 otherwise, yielding a bipolar one-hot-like, zero-centered encoding that improves training stability. The diffusion timestep *t* ∈ {1, …, *T*} with *T* = 200 is embedded into a 200-dimensional vector through a sinusoidal positional representation followed by a two-layer MLP with GELU activation. The condition labels, including MRL and MFE, are embedded into a 200-dimensional vector via a two-layer MLP with SiLU activation. These embeddings modulate intermediate UNet features, allowing the model to predict the noise *ε*_*t*_ conditioned on targets and sampling applies the reverse denoising update for *T* steps to transform Gaussian noise into valid sequences.

**Figure 1.**
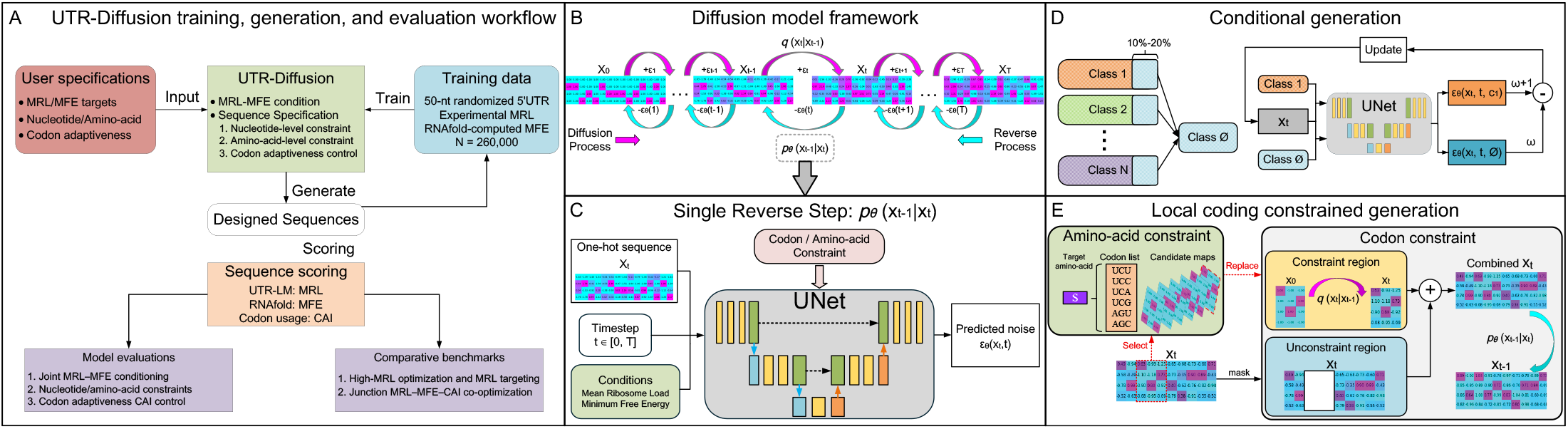
Overview of UTR-Diffusion for 5′ UTR and peptide-preserving 5′ UTR–CDS junction design. (A) Workflow and study design. UTR-Diffusion is trained on 50-nt randomized 5′ UTR sequences paired with experimentally measured MRL and RNAfold-computed MFE. At generation, users provide MRL–MFE targets, optional nucleotide or amino-acid specifications, and a codon-adaptiveness target. The generated designs are scored using UTR-LM for MRL, RNAfold for MFE, and relative codon-adaptiveness weights for CAI, and are analyzed in three model evaluations and two comparative benchmarks. (B) Diffusion model formulation. Gaussian noise is progressively added to one-hot-like sequence representations during the forward process and removed during reverse diffusion. (C) Single reverse-denoising step. At timestep *t*, the UNet receives the noisy sequence *x*_*t*_, the timestep embedding, and the MRL–MFE target embedding, and predicts the added noise. Sequence-control operations are applied during the reverse transition. (D) Classifier-free guidance with continuous-valued targets. During training, MRL–MFE target vectors are masked with probability 0.1–0.2, enabling the shared UNet to learn both conditional and masked-condition denoising. During sampling, predictions under the specified target vector and the masked condition are combined to strengthen adherence to the requested MRL–MFE targets. (E) Three modes of sequence control. Exact nucleotide subsequences are restored within constrained regions; amino-acid identities are preserved through dynamic selection among synonymous codons; and a codon-adaptiveness prior biases synonymous-codon selection to modulate sequence-level CAI.

### Classifier-Free Guidance

To strengthen conditional control and better align generated sequences with the target expression indicators, we adopt the classifier-free guidance (CFG) [Ho and Salimans, 2022]. During training, a portion (10%-20%) of the conditional labels are randomly masked, enabling the network to learn both conditional and unconditional denoising. As illustrated in Figure 1D, the network treats unconditional samples as an additional class and learns them jointly together with all the existing classes. At inference time, predictions from the unconditional branch *ε*_*θ*_ (*x*_*t*_, *t, ϕ*) and the conditional branch *ε*_*θ*_ (*x*_*t*_, *t, c*) are linearly combined to guide the reverse diffusion process toward the specified condition, as follows:

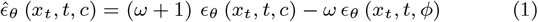

where the weight coefficient *ω* controls the guidance strength. The guided prediction 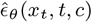 is then used to perform the reverse denoising step. This guidance explicitly steers the sampling trajectory toward the conditional signal while leveraging the unconditional estimate as a baseline.

### Three Forms of Sequence Specification

To combine MRL–MFE target control with user-defined sequence specifications, we adopt a RePaint-inspired resampling strategy [Lugmayr et al., 2022]. The resulting framework supports three complementary forms of specification: exact nucleotide-level constraints, amino-acid-level constraints with synonymous-codon flexibility, and codon-adaptiveness control over synonymous-codon selection.

#### Nucleotide-level constrained generation

Exact nucleotide-level control is used when a user-specified local sequence, such as a regulatory motif or an arbitrary nucleotide subsequence, must be strictly preserved. The specified subsequence is treated as a constrained region during sampling (Figure 1E). At each denoising step, the clean user-specified subsequence is forward-diffused to the current noise level, yielding the noisy representation of the constrained region. This constrained representation is then combined with the unconstrained region obtained from the preceding sampling state, and the assembled full sequence is updated by the denoising model. To maintain global coherence between constrained and unconstrained regions, noise is periodically re-injected into the sequence and followed by further denoising. This alternating re-noising procedure facilitates information propagation across the full sequence while strictly preserving the specified nucleotides.

#### Amino-acid-level constrained generation

Since the 5′ UTR and the proximal CDS jointly influence translation initiation and local RNA structure, 5′ UTR design should account for the downstream coding context rather than treating the two regions independently. We therefore introduce amino-acid-level constrained generation, which preserves specified amino-acid identities at the 5′ UTR–CDS junction while allowing synonymous codon variation. Because an amino-acid can generally be encoded by multiple synonymous codons, fixing a single codon would unnecessarily eliminate this sequence flexibility.

We enforce each amino acid as a set-valued nucleotide constraint whose codon realization is dynamically selected during sampling. For an amino-acid-constrained position *k* with target amino acid *a*, let *z*_*t,k*_ ∈ ℝ^4×3^ denote the current 3-nt representation at timestep *t*, and let 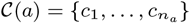 denote the *n*_*a*_ synonymous codons encoding *a*, each represented in the same one-hot-like 4×3 form. The candidate with the smallest squared Frobenius distance from the current 3-nt state is selected according to Equation 2:

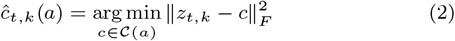

The selected codon 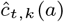 replaces the current 3-nt state at site *k*, after which sampling proceeds through the same RePaint-style denoising and periodic re-noising schedule used for nucleotide-level constrained generation. This procedure strictly preserves amino-acid identity while retaining synonymous-codon variability.

#### Codon-adaptiveness-controlled generation

Biasing synonymous-codon selection provides a route to control the expectation of relative codon adaptiveness at each amino-acid position and thereby indirectly modulate sequence-level CAI. For each codon *c*_*i*_ ∈ *C*(*a*), let *q*_*i*_ = *q*(*c*_*i*_ | *a*) denote its codon-usage frequency and let *w*_*i*_ = *q*_*i*_*/* max_*j*_ *q*_*j*_ denote its relative codon adaptiveness [Sharp and Li, 1987]. Given a user-specified adaptiveness target *α*, we seek a codon-selection prior *p*_*α*_(*c*_*i*_ | *a*) satisfying Equation 3:

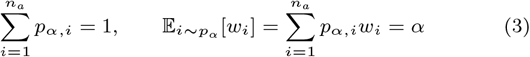

When an amino-acid has more than two synonymous codons, these constraints alone do not uniquely determine the probability vector. We therefore restrict the prior to a one-parameter exponential tilt of the natural codon-usage distribution:

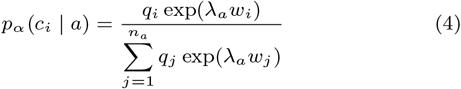

Because *q*_*i*_ and *w*_*i*_ are fixed, this reparameterization reduces moment matching in Equation 3 to a one-dimensional root-finding problem in *λ*_*a*_. For a feasible *α*, the resulting value of *λ*_*a*_ uniquely determines *p*_*α*_(· | *a*). Amino acids encoded by a single codon retain their unique codon without adaptiveness reweighting.

The resulting distribution is used as a codon-selection prior rather than sampled directly. At each denoising step, it reweights the squared Frobenius distance used for synonymous-codon selection:

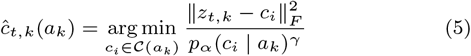

Here, *γ* controls the influence of the adaptiveness prior in each denoising-step codon replacement; repeated application of this rule biases the final synonymous-codon distribution over the reverse-diffusion trajectory. We used *γ* = 0.04, selected by the Supplementary sensitivity analysis, to enable indirect control of sequence-level CAI while preserving amino-acid identity without retraining the diffusion model.

### Dataset and Sequence Evaluation

We used 260,000 unique 50-nt randomized 5′ UTR sequences from the HEK293 MPRA library reported by [Sample et al., 2019]. Experimentally measured MRL and RNAfold-computed MFE [Hofacker, 2003] were used as the two continuous conditioning values during model training. Generated sequences were evaluated using a common sequence-scoring procedure. MRL was predicted using UTR-LM [Chu et al., 2024], MFE was computed using RNAfold, and CAI was calculated as the geometric mean of human relative codon adaptiveness values over CDS region [Sharp and Li, 1987].

Additional analyses included normalized MRL–MFE *L*_2_ error, pairwise Levenshtein distance, position-wise normalized Shannon entropy, GC content, and 3-mer motif richness. Detailed metric definitions, normalization procedures, and aggregation units are provided in the Supplementary Methods.

## Results

### Joint MRL–MFE Control in 5′ UTR Sequence Generation

We evaluated joint MRL–MFE control over a dense target grid by sweeping MRL from 4.0 to 8.0 in increments of 0.4 and MFE from −20 to 0 kcal mol^−1^ in increments of 2, yielding 121 joint target conditions. For each target pair, 100 sequences were generated and scored using UTR-LM and RNAfold.

At nine representative targets, the generated distributions were clearly separated, although their means showed target-dependent deviations from the requested coordinates (Figure 2A). Across the full grid, normalized *L*_2_ errors remained small over most of the target space but increased near the boundaries, particularly in the high-MRL and low-MFE corner; the mean error vectors generally pointed toward the interior (Figure 2B). Normalized pairwise Levenshtein distance and position-wise normalized Shannon entropy varied only modestly across the target grid and remained close to, but below, their random-sequence baselines, consistent with high yet structured sequence diversity (Figures 2D and E). Position-wise nucleotide profiles and the GC-content landscape showed increasing G/C enrichment as target MFE became more negative, broadly reproducing the corresponding trend in the training data (Figures 2C and F). The 3-mer motif-richness landscape also varied systematically across the target space and broadly followed the training-data contours, with larger deviations near the boundaries (Figure 2G).

**Figure 2.**
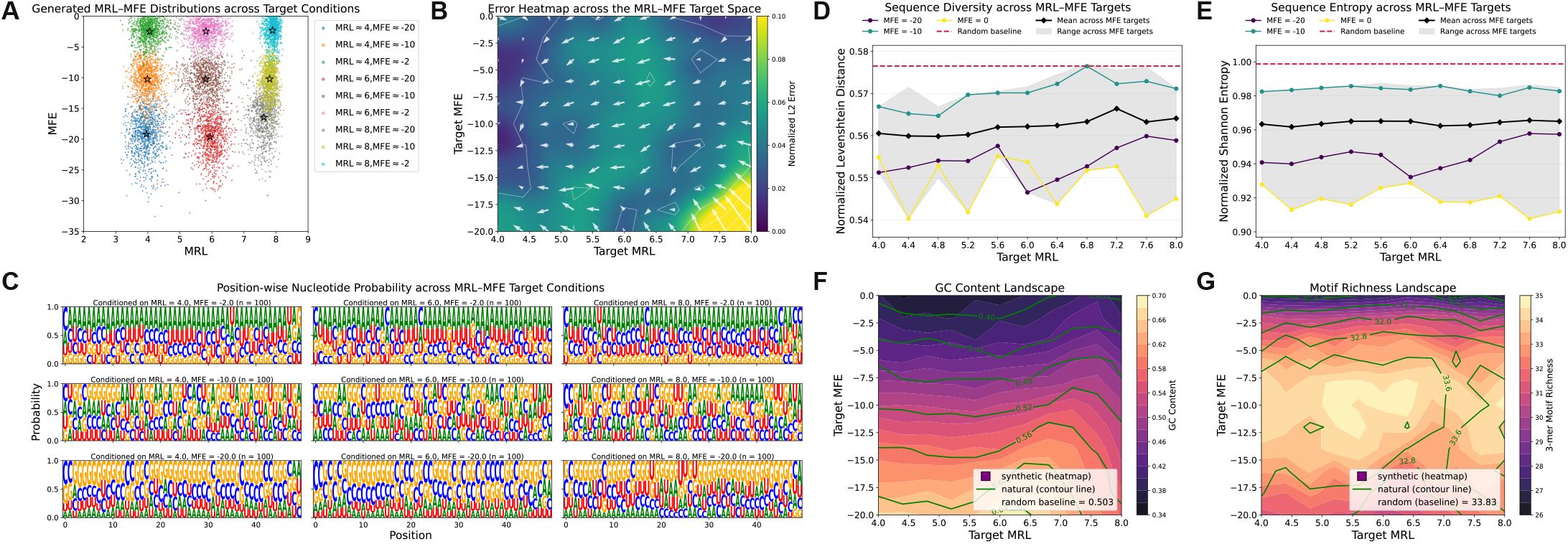
Joint MRL–MFE control across the target space. (A–B) Joint MRL–MFE response. Generated MRL–MFE distributions at nine representative target pairs; stars denote the mean of each generated distribution (A). Normalized L2 error landscape and mean error vectors across the full target grid; arrows denote the mean error vector at each requested target (B). (C) Target-dependent nucleotide composition. Mean position-wise nucleotide probabilities over *n* = 100 generated sequences for each representative target pair. (D–E) Sequence diversity. Normalized pairwise Levenshtein distance (D) and position-wise normalized Shannon entropy (E) across MRL targets for selected MFE conditions. Black lines denote the mean across MFE conditions, gray bands show the corresponding range, and dashed lines indicate random-sequence baselines. (F–G) Sequence composition. GC content (F) and 3-mer motif richness (G) of generated sequences shown as heatmaps across the MRL–MFE target space, with training-data contours and random-sequence baselines for comparison.

Overall, UTR-Diffusion responded jointly to MRL and MFE targets while maintaining high sequence diversity and broadly preserving training-data compositional patterns.

### Joint MRL–MFE Control under Nucleotide- and Amino-Acid-Level Constraints

We next examined whether joint MRL–MFE control was maintained under explicit nucleotide- and amino-acid-level constraints. The two constraint modes were evaluated over the same MRL–MFE target grid used in the preceding evaluation (Figures 3A and B). The nucleotide-level series comprised four pattern families constructed from Kozak, DRACH, and randomly selected triplet variants, whereas the amino-acid-level series retained a fixed AUG and constrained one to four downstream amino-acid sites. Each constraint family comprised multiple variants; their allowed identities and complete definitions are provided in Supplementary Table S2.

**Figure 3.**
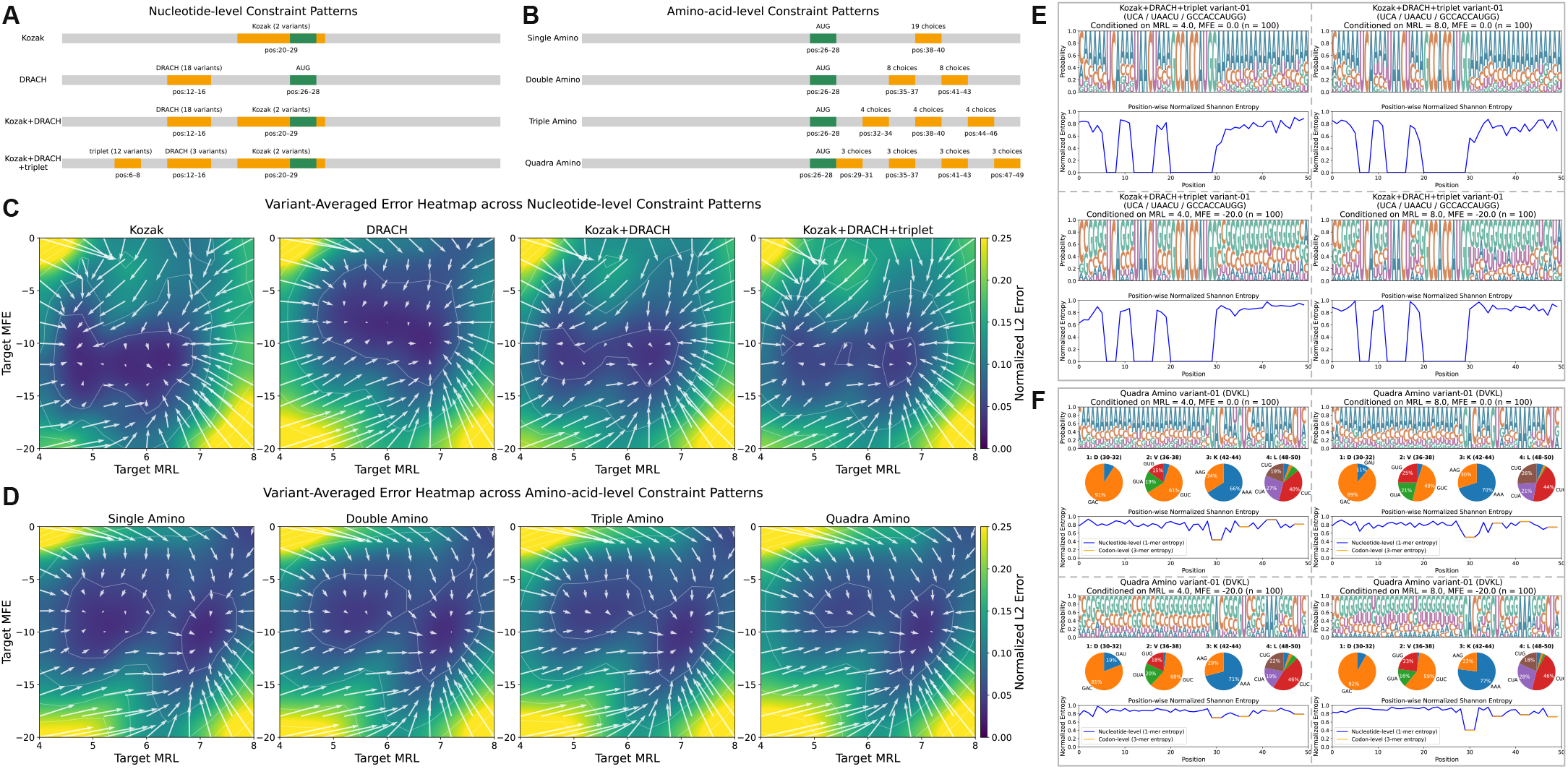
Joint MRL–MFE control under nucleotide- and amino-acid-level constraints. (A–B) Constraint designs. Four nucleotide-level constraint families constructed from Kozak, DRACH, and randomly selected triplet variants (A), and four amino-acid-level constraint families containing a fixed AUG and one to four downstream amino-acid sites (B). Each family comprises multiple variants whose allowed identities are listed in Supplementary Table S2. (C–D) Variant-averaged MRL–MFE target errors. Normalized *L*_2_ error landscapes and mean error vectors across the MRL–MFE target grid for the nucleotide-level (C) and amino-acid-level (D) families. Each heatmap cell and vector summarize the generated-mean error averaged across all variants in the corresponding family at that target; color and arrow length indicate error magnitude, and arrow direction indicates the direction of the mean target error. (E–F) Constraint preservation and local diversity. Representative variant-01 generations at four corner targets (MRL ∈ {4, 8}; MFE ∈{0, −20} kcal mol^−1^; *n* = 100 per target) for the Kozak+DRACH+triplet family (E) and quadruple-amino family (F). Position-wise nucleotide probabilities and normalized base- and triplet-level Shannon entropy are shown for both families; synonymous-codon usage at the four downstream amino-acid-constrained sites is additionally summarized in (F).

Relative to unconstrained generation (Figure 2B), the normalized MRL–MFE *L*_2_ errors increased modestly, with the clearest increases near the boundaries of the MRL–MFE target space (Figures 3C and D). Within each constraint mode, errors also tended to increase slightly as the nucleotide-level patterns became more complex or additional amino-acid sites were constrained. Nevertheless, broad low-error regions remained over the central target space, indicating that joint MRL–MFE control was largely retained under both constraint modes.

To illustrate constraint enforcement and local sequence diversity, we selected variant 01 from the Kozak+DRACH+triplet and quadruple-amino families at the four corner targets defined by MRL ∈ {4, 8} and MFE ∈ {0, −20} kcal mol^−1^ (Figures 3E and F). Under nucleotide-level constraints, the specified Kozak, DRACH, and triplet sequences were preserved at their intended positions, while unconstrained sites retained substantial positional diversity and target-dependent nucleotide patterns broadly similar to those observed without local constraints. Under amino-acid-level constraints, the fixed AUG and specified downstream amino-acid identities were preserved, multiple synonymous codons remained represented at the constrained sites, and unconstrained positions retained substantial nucleotide diversity. Together, these results show that nucleotide- and amino-acid-level constraints can be integrated with joint MRL–MFE control with only a modest increase in target error, while preserving diversity outside the constrained regions and synonymous-codon flexibility at amino-acid-constrained sites.

### Joint MRL–MFE and Codon-Adaptiveness Control at the 5′ UTR–CDS Junction

We then evaluated joint MRL–MFE and codon-adaptiveness control while preserving predefined downstream peptide sequences. Each 50-nt design comprised a 26-nt 5′ UTR without position-specific constraints, a fixed AUG, and seven downstream codons encoding one of 10 reference peptides (Supplementary Table S3). MRL targets {4, 6, 8} and MFE targets {−20, −10, −2} kcal mol^−1^ were combined with adaptiveness targets *α* ∈ {0.85, 0.90, 0.95}; 100 sequences were generated for each peptide–MRL–MFE–*α* combination.

The 30 peptide–adaptiveness group means shifted systematically across the nine requested MRL–MFE conditions (Figure 4A). The three MFE response levels remained clearly separated, although generated values were shifted toward more negative MFE at the less-negative targets. By contrast, generated MRL values were compressed toward the middle of the requested range, resulting in a more pronounced loss of target resolution along the MRL axis. Within each target panel, the three *α* groups largely overlapped, whereas points carrying the same peptide identifier often lay close to one another across the three adaptiveness settings. This pattern suggests that the constrained peptide context contributed more strongly than the tested adaptiveness range to within-target variation in generated MRL and MFE.

**Figure 4.**
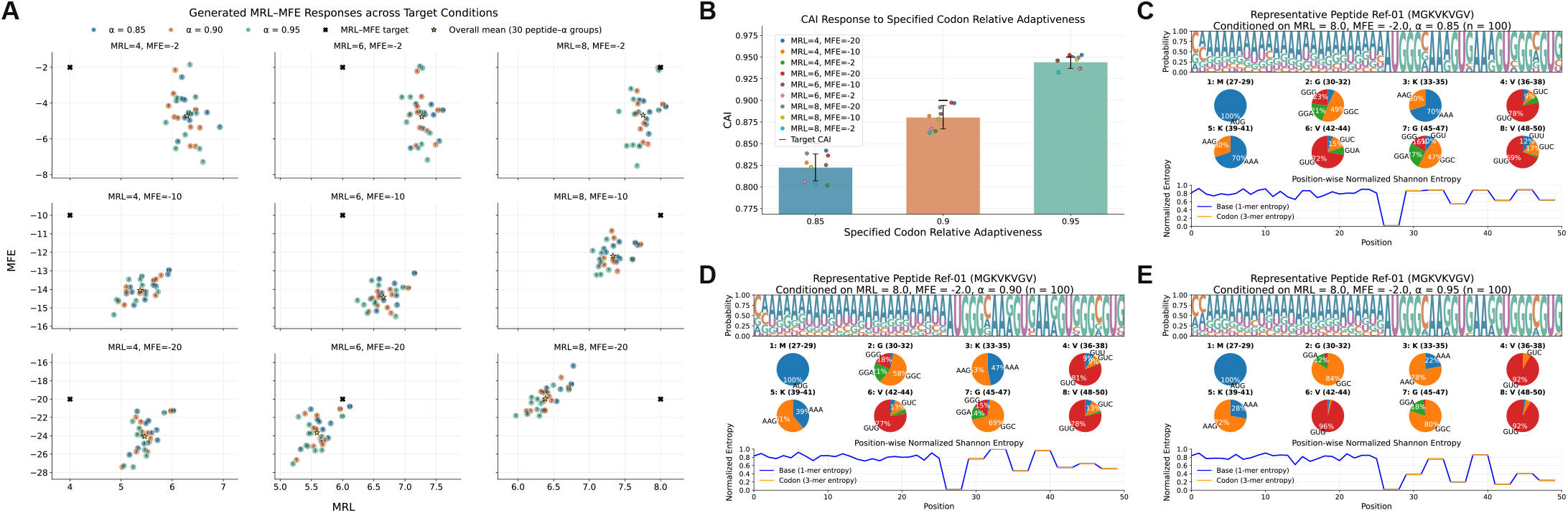
Joint MRL–MFE and codon-adaptiveness control under fixed peptide constraints at the 5′ UTR–CDS junction. (A) Generated MRL–MFE responses across nine target pairs. Each colored, numbered point represents the mean of *n* = 100 sequences generated for one reference peptide and one adaptiveness target. Color denotes *α* = 0.85, 0.90, or 0.95, and the numeral identifies one of the 10 reference peptides listed in Supplementary Table S3. Points carrying the same numeral therefore represent the same peptide under the three adaptiveness settings. Black crosses indicate the requested MRL–MFE targets, and gold stars denote the mean across all 30 peptide–adaptiveness groups in each target panel. (B) Sequence-level CAI response to the specified codon-adaptiveness target. Each colored point represents the mean across the 10 reference peptides for one MRL–MFE target condition. Bars and error bars show the mean ± SD across the nine target-condition means, and black horizontal markers indicate *α* = 0.85, 0.90, and 0.95. CAI was calculated over the seven downstream codons. (C–E) Representative generations for Ref-01 (MGKVKVGV) at MRL = 8, MFE = −2 kcal mol^−1^, and *α* = 0.85 (C), 0.90 (D), or 0.95 (E), with *n* = 100 sequences per condition. Each panel shows position-wise nucleotide probabilities, codon-usage distributions at the fixed AUG and seven downstream peptide positions, and normalized base- and codon-level Shannon entropy. Increasing *α* shifts synonymous-codon usage toward higher-adaptiveness codons, while the encoded peptide is preserved and the 26-nt 5′ UTR retains substantial nucleotide diversity.

Generated sequence-level CAI increased monotonically with the specified codon-adaptiveness target (Figure 4B). Mean CAI values were approximately 0.82, 0.88, and 0.94 for *α* = 0.85, 0.90, and 0.95, respectively, with limited variation across the nine MRL–MFE conditions. The small offsets between CAI and *α* are consistent with their different definitions: *α* specifies the position-level expectation of relative codon adaptiveness, whereas each generated sequence realizes seven discrete codon weights whose geometric mean defines CAI.

For the representative peptide Ref-01 (MGKVKVGV), we further compared generations at MRL = 8, MFE = −2, and the three adaptiveness targets (Figures 4C–E). As *α* increased, synonymous-codon usage shifted toward codons with higher relative adaptiveness and became more concentrated at several peptide positions, while the 26-nt 5′ UTR retained high position-wise nucleotide entropy. Together, these results show that UTR-Diffusion integrates joint MRL–MFE conditioning, peptide preservation, and graded codon-adaptiveness control while retaining substantial 5′ UTR sequence diversity.

### High-MRL Optimization and Precise MRL Targeting in 5′ UTR Design

We benchmarked UTR-Diffusion in two 50-nt 5′ UTR design tasks: high-MRL optimization and precise MRL targeting.

In the high-MRL optimization task (Figure 5A), UTR-Diffusion was compared with UTRGAN [Barazandeh et al., 2025] and Optimus 5-Prime [Sample et al., 2019], with 1,000 sequences generated for each method. Optimus and UTR-Diffusion were conditioned on MRL = 9, an extrapolative target, whereas UTRGAN was evaluated using its standard high-score generation procedure. UTRGAN outputs longer than 100 nt were excluded prior to evaluation, and a random top 1% reference was derived from 100,000 random 50-nt sequences. UTR-Diffusion achieved the highest mean MRL with a compact distribution, exceeding Optimus and the random top 1% reference, while UTRGAN showed a lower mean and substantially greater variability.

**Figure 5.**
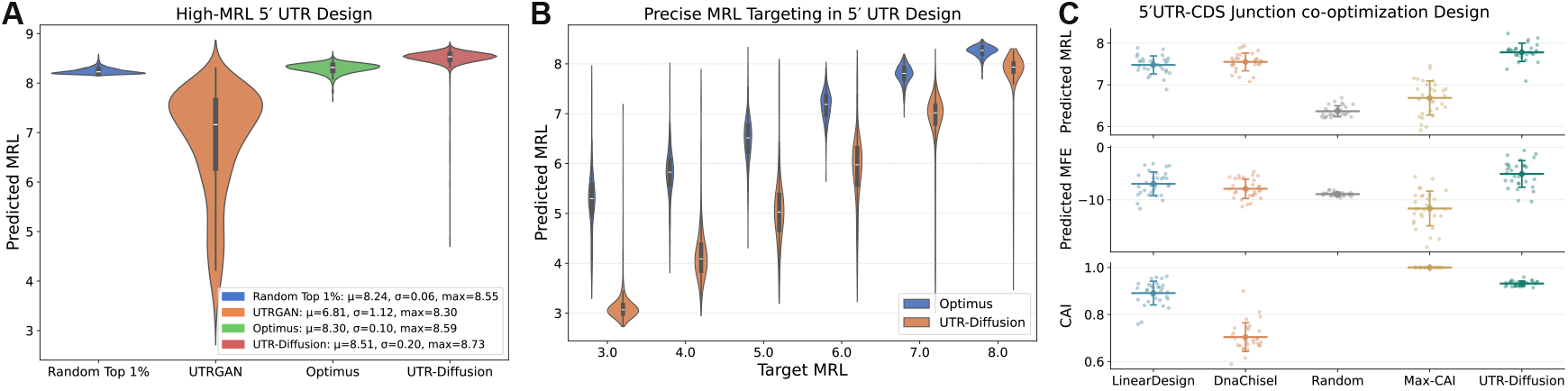
Benchmarking UTR-Diffusion against existing sequence design methods in 5′ UTR and 5′ UTR–CDS junction design. (A) High-MRL 5′ UTR design: distributions of predicted MRL over *N* = 1,000 sequences per method, including a random top 1% reference baseline; UTRGAN sequences longer than 100 nt were excluded prior to evaluation. (B) Precise MRL targeting in 5′ UTR design: predicted MRL distributions for Optimus and UTR-Diffusion across requested targets (MRL = 3–8). (C) Multi-objective 50-nt 5′ UTR–CDS junction design evaluated by predicted MRL (top), MFE (middle), and CAI (bottom). A higher, less-negative MFE indicates lower predicted folding propensity within the local junction. UTR-Diffusion (*α* = 0.95) is compared with LinearDesign, DNAChisel, an unconstrained random control, and a single-objective Max-CAI reference. Each small point represents the mean of 100 candidates generated for one of 30 reference peptides; large markers and error bars show the mean ± SD across the 30 reference-level means. CAI was calculated over the nine codons downstream of the fixed AUG and was not reported for the unconstrained random control.

In the precise MRL-targeting task (Figure 5B), UTR-Diffusion and Optimus were evaluated at target values {3, 4, 5, 6, 7, 8}, with 1,000 sequences generated for each method at each target. UTR-Diffusion closely tracked the requested values with relatively narrow distributions, whereas Optimus produced broader distributions and a systematic upward shift, most prominently at low and intermediate targets.

Together, UTR-Diffusion achieved the strongest performance in both benchmarks, combining the highest mean MRL in the optimization task with substantially more accurate MRL target tracking.

### Multi-objective Optimization of MRL, MFE, and CAI in 5′ UTR–CDS Junction Design

We next benchmarked UTR-Diffusion in a 50-nt 5′ UTR–CDS junction design task requiring simultaneous optimization of MRL, MFE, and CAI. Because reduced folding around the 5′ leader and start-codon-proximal CDS can improve translation-initiation accessibility[Kozak, 1989, Liebhaber et al., 1992, Mauger et al., 2019], the benchmark jointly targeted high MRL, high (less-negative) junction MFE, and high CAI.

Each design comprised a 20-nt 5′ UTR, a fixed AUG, and a 27-nt synonymous CDS segment encoding one of 30 nine-amino-acid reference peptides derived from human proteins (Supplementary Table S4). For each reference peptide, every method generated 100 candidates while preserving the nine downstream amino-acid identities. LinearDesign [Zhang et al., 2023] and DNAChisel [Zulkower and Rosser, 2020] were used as peptide-preserving baselines, with LinearDesign applying its 5′-leader opening heuristic and DNAChisel combining AvoidHairpins with MaximizeCAI; an unconstrained random control and a single-objective Max-CAI reference were also included. UTR-Diffusion was conditioned on MRL = 9, MFE = 0 kcal mol^−1^, and the highest specified codon-adaptiveness target, *α* = 0.95.

UTR-Diffusion achieved the highest mean MRL and MFE among all evaluated groups and a higher CAI than both peptide-preserving baselines (Figure 5C). The random control showed lower MRL and MFE, whereas Max-CAI reached CAI = 1 by construction but at the cost of lower MRL and more negative junction MFE. Overall, UTR-Diffusion provided the strongest balance of high MRL, high junction MFE, and high CAI while preserving the encoded peptide sequence.

## Discussion

### Integrated Control of Quantitative Targets and Sequence Specifications

In this study, we developed a diffusion-based framework for 5′ UTR and 5′ UTR–CDS junction design that combines numerical targets for MRL and MFE with three forms of sequence specification: exact nucleotide constraints, amino-acid identity constraints with synonymous-codon flexibility, and codon-adaptiveness control for graded modulation of sequence-level CAI. Across the three model evaluations, UTR-Diffusion retained joint MRL–MFE responsiveness while preserving specified nucleotide subsequences or peptide identities, and increasing the codon-adaptiveness target produced monotonic shifts in CAI. The external benchmarks further showed accurate MRL targeting in 5′ UTR design and a favorable balance of high MRL, high junction MFE, and high CAI in peptide-preserving junction design.

The central contribution is the integration of heterogeneous design specifications within a single generative procedure. MRL and MFE are supplied as numerical target values; exact nucleotide identities are imposed through RePaint-style constraint enforcement; amino-acid identities are represented by sets of synonymous codons; and codon adaptiveness biases selection among those codons. This organization avoids decomposing the design problem into separate stages whose outputs must subsequently be reconciled. The modest increase in MRL–MFE error as more positions or coding identities were specified reflects the expected trade-off between user-imposed specifications and the sequence freedom remaining to satisfy the numerical targets.

With suitable training labels or validated scoring models, the same target-conditioning mechanism could in principle be extended to other quantitative sequence properties, such as local RNA accessibility or transcript half-life. Similarly, new sequence specifications could be implemented by fixing exact nucleotides, defining sets of admissible sequence realizations, or assigning graded preferences among admissible choices—the same mechanisms used here for motifs, synonymous codons, and codon adaptiveness, respectively. This combination of numerical target control and flexible sequence specification provides a general strategy for multi-objective RNA design, although achievable performance will remain bounded by training-data coverage, scoring-model accuracy, and the sequence degrees of freedom remaining after user specifications are imposed.

### Boundary Behavior and Limited Extrapolation in the MRL–MFE Landscape

UTR-Diffusion exhibited its most informative behavior near the boundaries of the MRL–MFE target space. Within well-supported regions, the generated means generally followed the requested targets, whereas boundary conditions produced larger errors and systematic displacement toward the interior of the evaluated region. In the high-MRL benchmark, individual generated sequences nevertheless reached predicted MRL values beyond the upper range observed for training sequences scored using the same predictor. This combination of inward-biased distribution means and extreme tail samples supports limited extrapolative generation, but not uniformly calibrated control outside the training-supported region. Target adherence therefore appears to depend not only on the conditioning signal, but also on the availability of compatible sequences in the underlying training distribution.

The high-MRL and strongly negative-MFE corner was particularly difficult to realize. This pattern is consistent with the biological tension between stable 5′ UTR secondary structure and efficient translation initiation, because extensive base pairing can impede ribosomal scanning and start-codon accessibility [Kozak, 1989, Liebhaber et al., 1992, Mauger et al., 2019]. At the same time, generated sequences retained high Levenshtein distance and nucleotide entropy within individual target conditions, indicating that achievable regions of the MRL–MFE landscape admit multiple distinct sequence solutions rather than a single dominant design. Thus, the observed target-space behavior reflects a trade-off between translation-related output and predicted RNA folding propensity, while also revealing substantial sequence degeneracy within feasible regions.

### Limitations of the Present Study

The present study has three principal limitations.

First, the model was trained on a randomized 5′ UTR MPRA performed in HEK293 cells. These laboratory-synthesized sequences and reporter constructs may differ from endogenous transcripts in sequence composition and regulatory context. Moreover, the experimental MRL values were measured within a shared reporter context, and the learned sequence–MRL relationships may therefore partly depend on this assay architecture. Generalization to other cell types, reporter constructs, or endogenous transcripts remains to be evaluated and may require retraining or fine-tuning on data from the relevant biological context.

Second, the current UNet implementation operates on fixed-length 50-nt representations and therefore does not natively support variable-length or full-length mRNA generation. Extending the framework to longer sequences would require architectural and training modifications, together with sufficiently large and well-annotated long-sequence datasets, while full-resolution diffusion would incur substantially greater computational cost. Latent diffusion approaches [Huang et al., 2024, Li et al., 2024] may reduce this cost by operating in compressed representations, but preserving exact nucleotide and amino-acid specifications through latent encoding and decoding would require additional methodology. Accordingly, the present results characterize local 5′ UTR and 5′ UTR–CDS junction design rather than full-length transcript optimization.

Finally, newly generated sequences were evaluated computationally rather than experimentally. MRL was assessed using UTR-LM, whereas MFE was defined and evaluated using RNAfold. Although UTR-LM predictions correlate with experimentally measured MRL (Supplementary Figure S11) and RNAfold provides a widely used estimate of RNA folding propensity, these scores do not establish protein output or RNA structure in cells. In particular, because RNAfold was used for both MFE labeling and evaluation, the reported MFE control should be interpreted relative to the RNAfold-defined metric. We did not perform reporter assays, MPRA, or direct structural measurements on the generated sequences; experimental validation is therefore required before practical application of the proposed designs.

## Conclusions

In summary, UTR-Diffusion provides a unified generative framework for 5′ UTR and 5′ UTR–CDS junction design by combining numerical MRL–MFE targets with exact nucleotide constraints, amino-acid preservation, and codon-adaptiveness control. Across the model evaluations, the framework preserved specified nucleotide or peptide identities, enabled graded modulation of sequence-level CAI, and retained substantial sequence diversity. External benchmarks further demonstrated strong high-MRL optimization, accurate MRL target tracking, and a favorable balance of MRL, junction MFE, and CAI under peptide-preserving junction design. With experimental validation and iterative model–experiment feedback, this framework could provide a practical basis for multi-objective mRNA sequence design.

