## Supplementary Material for "UTR-Diffusion: Conditional Diffusion Modeling for Multi-objective and Constrained UTR Design"

### S1 Supplementary Figures

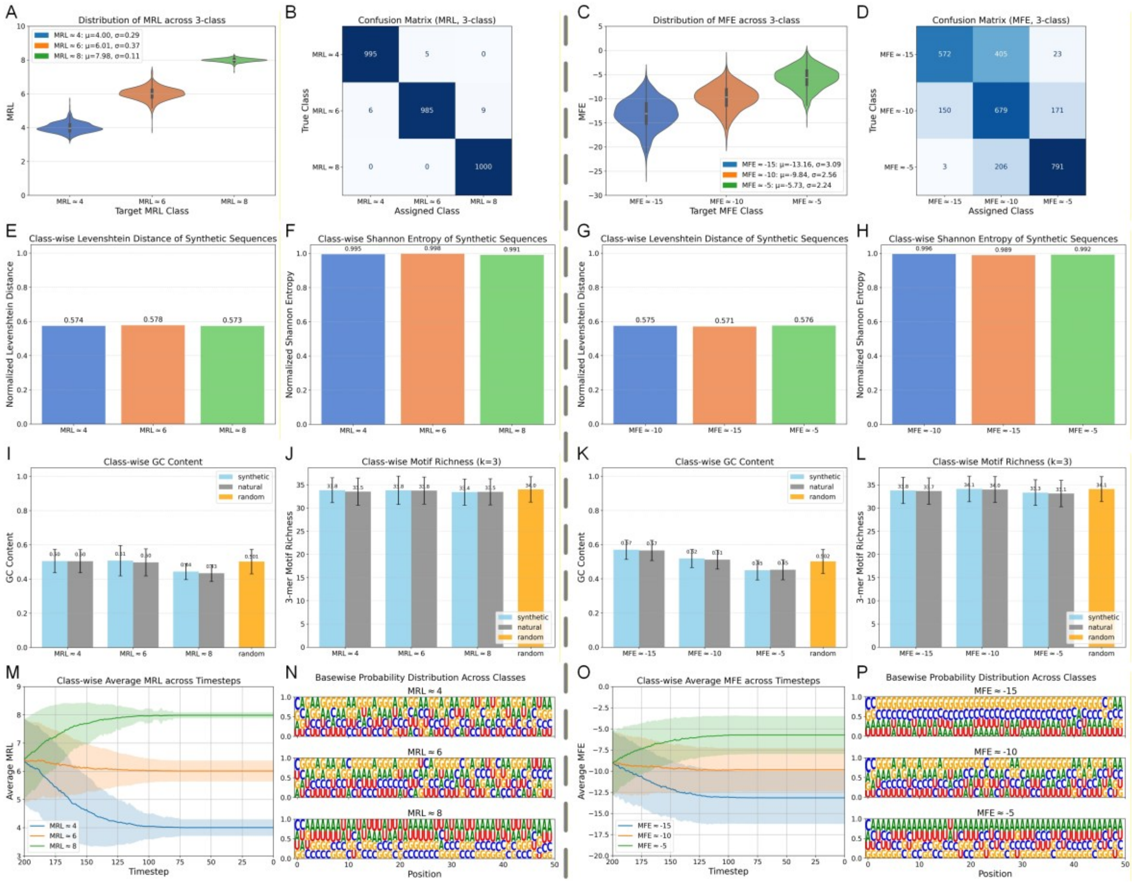

**Figure S 1: Discrete single-label conditional generation for MRL and MFE**

Conditional fidelity, generative diversity, and sequence composition under discrete single-indicator conditioning are evaluated for MRL (left block) and MFE (right block).

(A–D) Conditional fidelity. Violin plots show the distribution of predicted MRL (A) and MFE (C) across three discrete target classes, with corresponding confusion matrices (B, D) quantifying class-level assignment accuracy.

(E–H) Generative diversity. Class-wise normalized Levenshtein distance (E, G) and position-wise normalized Shannon entropy (F, H) assess intra-class sequence diversity under each conditioning setting.

(I–L) Compositional diagnostics. GC content (I, K) and 3-mer motif richness (J, L) of synthetic sequences are compared with natural training sequences and random controls.

(M, O) Indicator dynamic. Indicator trajectories illustrate the convergence of predicted indicator values toward their respective targets over sampling timesteps of diffusion reverse process.

(N, P) Sequence-level patterns. Class-wise nucleotide probability logos visualize position-specific nucleotide preferences under each MRL or MFE condition.

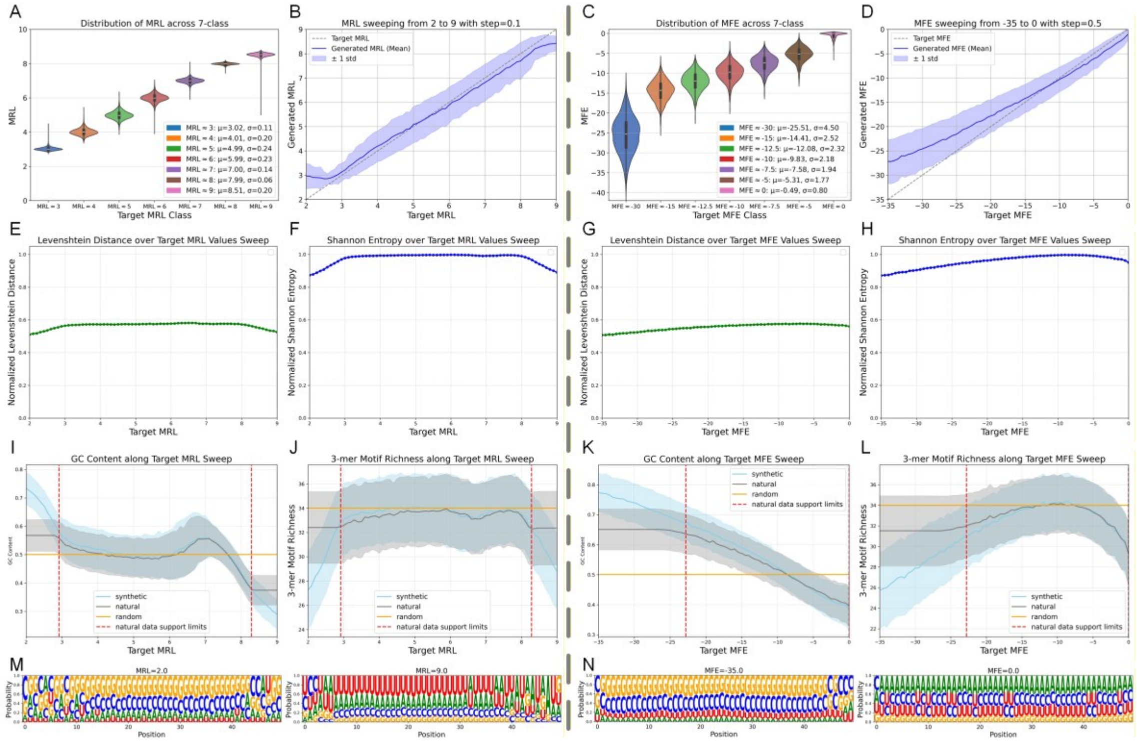

**Figure S 2: Continuous single-label conditional generation for MRL and MFE**

Model behavior under continuous target sweeps is evaluated for MRL (left block) and MFE (right block).

(A–D) Conditional fidelity. Violin plots summarize predicted MRL (A) and MFE (C) across seven representative target values sampled from dense sweeps (MRL 2–9; MFE −35–0). Sweep curves (B, D) show the mean ±1 std of generated indicator values as a function of the target, demonstrating near-linear tracking across the full range.

(E–H) Generative diversity. Normalized Levenshtein distance (E, G) and position-wise normalized Shannon entropy (F, H) remain high and stable across target sweeps, indicating preserved sequence-level and nucleotide-level variability.

(I–L) Compositional diagnostics. GC content (I, K) and 3-mer motif richness (J, L) of synthetic sequences co-vary with targets in close alignment with natural data, while remaining distinct from random controls.

(M–N) Sequence-level patterns. Representative nucleotide probability logos at extreme targets illustrate systematic positional reshaping of sequence composition along the continuous label axis.

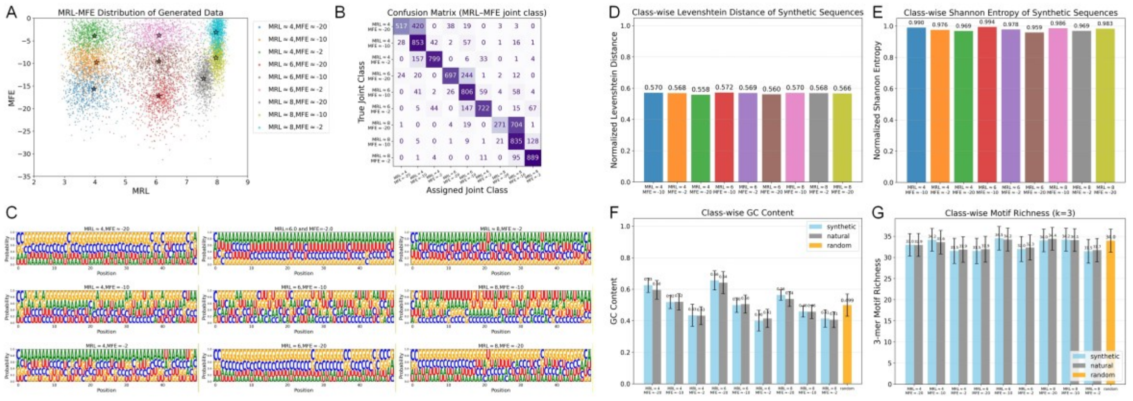

**Figure S 3: Discrete multi-label conditional generation for joint MRL–MFE control**

Joint conditional behavior under discrete multi-label settings is evaluated across the MRL–MFE plane.

(A–B) Conditional fidelity. Joint distributions of predicted MRL and MFE form well-separated clusters corresponding to the nine discrete target combinations (A). The confusion matrix (B) quantifies class-wise assignment consistency under joint conditioning.

(C) Sequence-level patterns. Class-specific nucleotide probability logos illustrate distinct positional base preferences across different MRL–MFE combinations.

(D–E) Generative diversity. Class-wise normalized Levenshtein distance (D) and normalized Shannon entropy (E) remain high across joint conditions, indicating preserved sequence-level and nucleotide-level variability under coupled control.

(F–G) Compositional diagnostics. GC content (F) and 3-mer motif richness (G) of synthetic sequences are compared with natural training data and random controls, demonstrating alignment with natural distributions across joint classes.

Together, these analyses indicate that the diffusion model enables coherent two-dimensional control of MRL and MFE while maintaining generative diversity and natural-like compositional characteristics.

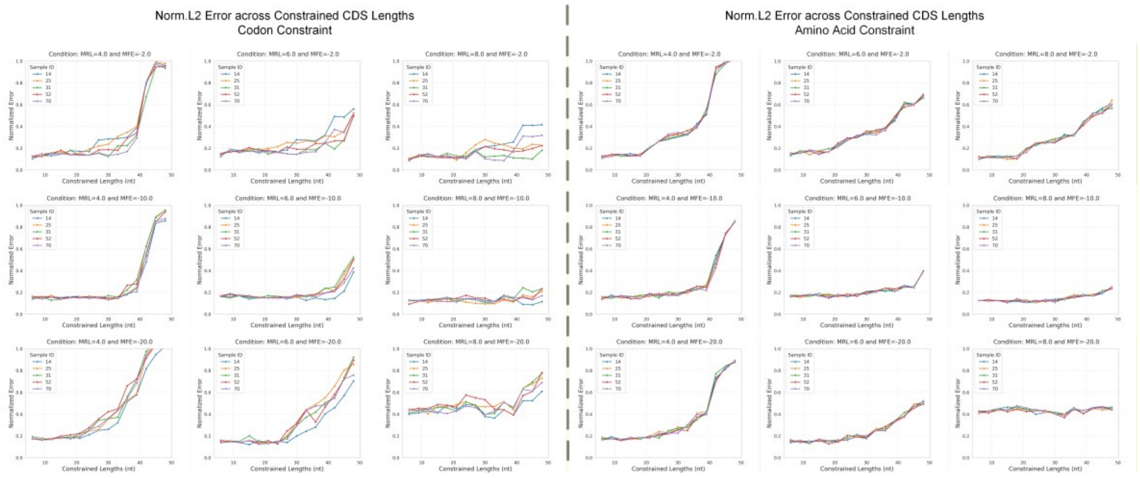

Figure S 4: Normalized L2 error under increasing constrained CDS lengths for codon- and amino-acid-level constraints

Normalized L2 error between generated sequences and the target conditions is plotted as a function of constrained CDS length (6–48-nt). The left block shows results under codon-level constraints, while the right block shows results under amino-acid-level constraints. Each block consists of nine subpanels, corresponding to nine distinct joint target settings of MRL and MFE. Within each subpanel, the x-axis denotes the length of the constrained CDS region, and the y-axis denotes the normalized L2 error between the generated sequences and the specified target conditions. Multiple curves represent the five predefined CDS-like reference segments under the same target setting.

Across all target settings and both constraint types, conditional error generally increases as the constrained CDS length grows, indicating progressively reduced freedom for satisfying global conditions under stronger local constraints. Notably, the rate of error increase varies substantially across different joint target pairs, suggesting that conditional controllability depends on the specific combination of MRL and MFE targets.

Despite these differences, codon-level and amino-acid-level constraints exhibit largely similar trends for corresponding target settings, with a pronounced degradation in conditional fidelity observed when the constrained region exceeds approximately 40-nt. This behavior suggests a shared upper limit of conditional controllability under strong local constraints, largely independent of the representation level of the imposed constraints.

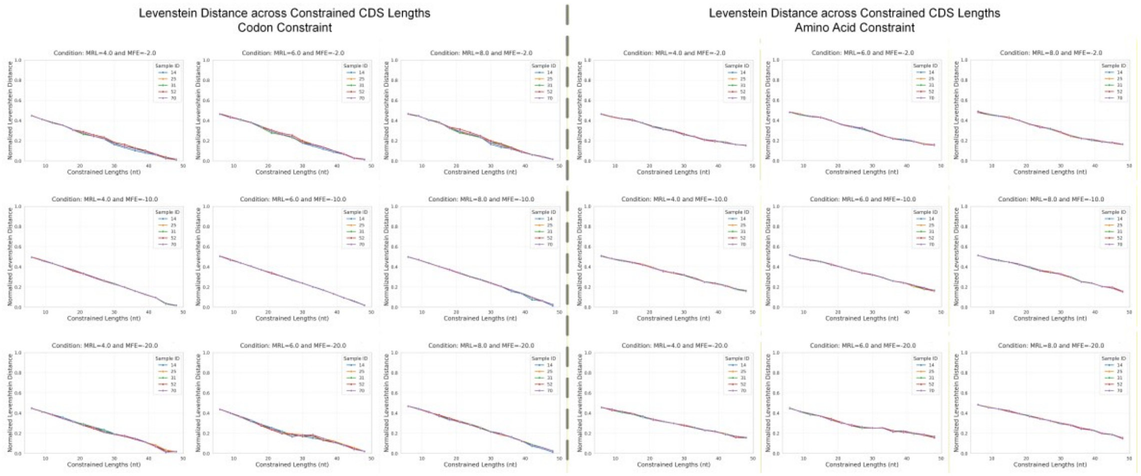

Figure S 5: Normalized Levenshtein distance under increasing constrained CDS lengths for codon- and amino-acid-level constraints

Normalized Levenshtein distance between generated sequences and reference sequences is plotted as a function of constrained CDS length (6–48-nt). The left block shows results under codon-level constraints, while the right block shows results under amino-acid-level constraints.

Each block consists of nine subpanels, corresponding to nine distinct joint target settings of MRL and MFE. Within each subpanel, the x-axis denotes the length of the constrained CDS region, and the y-axis denotes the normalized Levenshtein distance between generated and reference sequences. Multiple curves represent the five predefined CDS-like reference segments under the same target setting.

Across all target settings and both constraint types, the normalized Levenshtein distance decreases monotonically as the constrained CDS length increases, reflecting progressively stronger sequence similarity enforced by longer locally constrained regions. The overall trends are highly consistent across different joint target pairs and between codon- and amino-acid-level constraints, indicating that local constraint strength, rather than the representation level of the constraint, dominates the degree of sequence convergence.

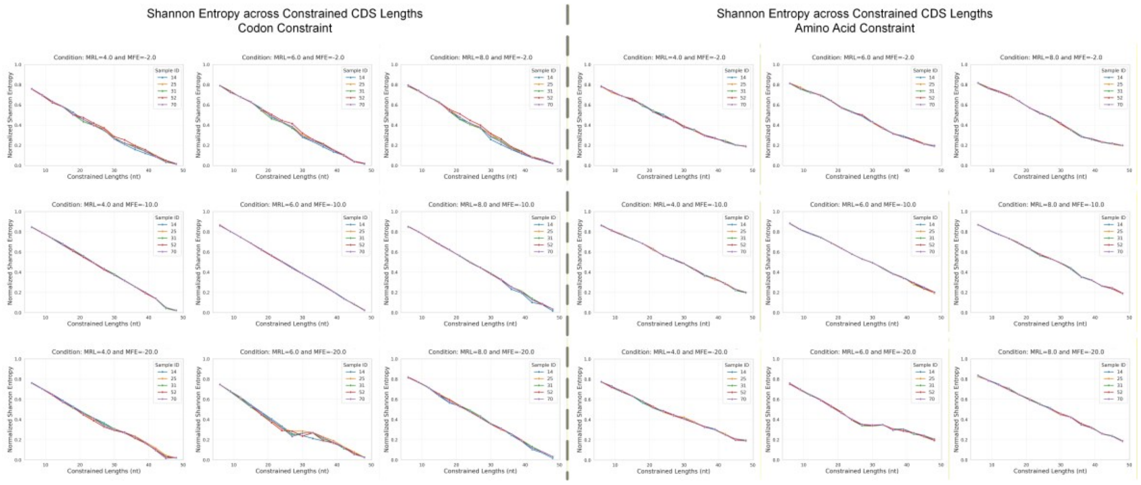

Figure S 6: Normalized Shannon entropy under increasing constrained CDS lengths for codon- and amino-acid-level constraints

Normalized Shannon entropy of generated sequences is plotted as a function of constrained CDS length (6–48-nt). The left block shows results under codon-level constraints, while the right block shows results under amino-acid-level constraints.

Each block consists of nine subpanels, corresponding to nine distinct joint target settings of MRL and MFE. Within each subpanel, the x-axis denotes the length of the constrained CDS region, and the y-axis denotes the normalized Shannon entropy of the generated sequences. Multiple curves represent the five predefined CDS-like reference segments under the same target setting.

Across all target settings and both constraint types, Shannon entropy decreases monotonically as the constrained CDS length increases, indicating a progressive reduction in sequence diversity induced by stronger local constraints. The observed trends are highly consistent across different joint target pairs and between codon- and amino-acid-level constraints, suggesting that constraint strength, rather than constraint representation, primarily governs the loss of sequence diversity.

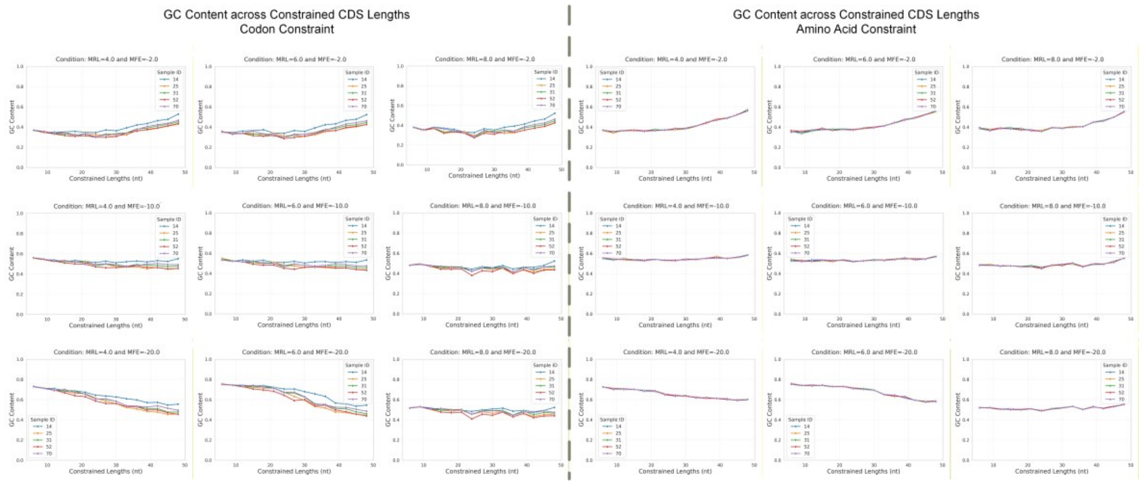

Figure S 7: GC content under increasing constrained CDS lengths under codon- and amino-acid-level constraints

GC content of generated sequences is plotted as a function of constrained CDS length (6–48-nt). The left block shows results under codon-level constraints, while the right block shows results under amino-acid-level constraints.

Each block consists of nine subpanels, corresponding to nine distinct joint target settings of MRL and MFE. Within each subpanel, the x-axis denotes the length of the constrained CDS region, and the y-axis denotes the GC content of the generated sequences. Multiple curves represent the five predefined CDS-like reference segments under the same target setting.

Across all target settings and both constraint types, GC content remains within a relatively narrow range as the constrained CDS length increases. While modest, target-dependent GC shifts are observed in some codon-constrained settings, no systematic monotonic trend is present. These results indicate that the changes in conditional fidelity, sequence similarity, and diversity reported in Supplementary Figures S 4 and S 5 are not primarily driven by global base-composition biases.

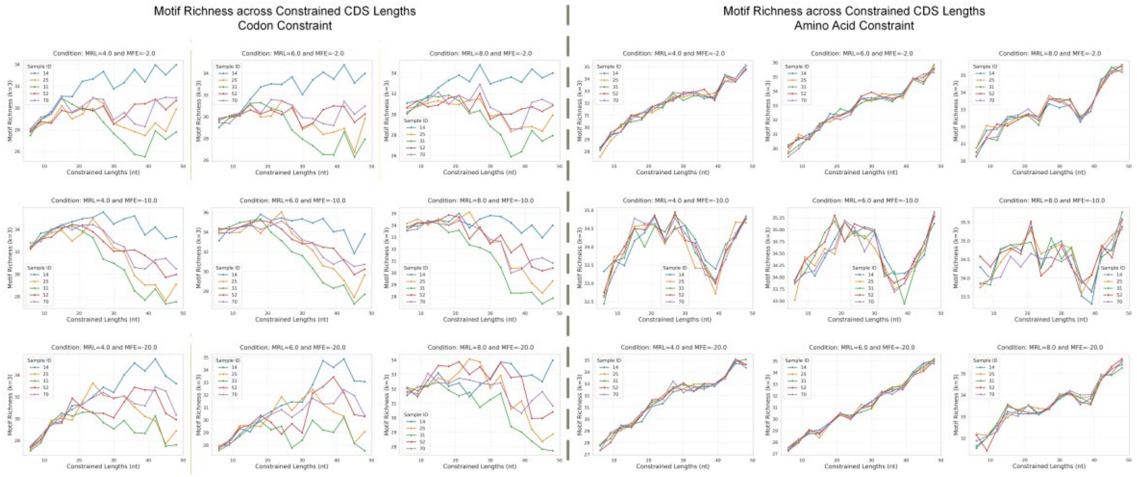

Figure S 8: Motif richness under increasing constrained lengths under codon- and amino-acid-level constraints

Motif richness (k = 3) of generated sequences is plotted as a function of constrained CDS length (6–48-nt). The left block shows results under codon-level constraints, while the right block shows results under amino-acid-level constraints.

Each block consists of nine subpanels, corresponding to nine distinct joint target settings of MRL and MFE. Within each subpanel, the x-axis denotes the length of the constrained CDS region, and the y-axis denotes the motif richness of the generated sequences. Multiple curves represent the five predefined CDS-like reference segments under the same target setting.

Across all target settings, increasing constraint length progressively limits local sequence flexibility, leading to a partial reduction in motif richness. However, substantial motif diversity is retained across constrained lengths, particularly under amino-acid-level constraints, indicating that higher-level constraints preserve combinatorial sequence variability even under strong local restrictions.

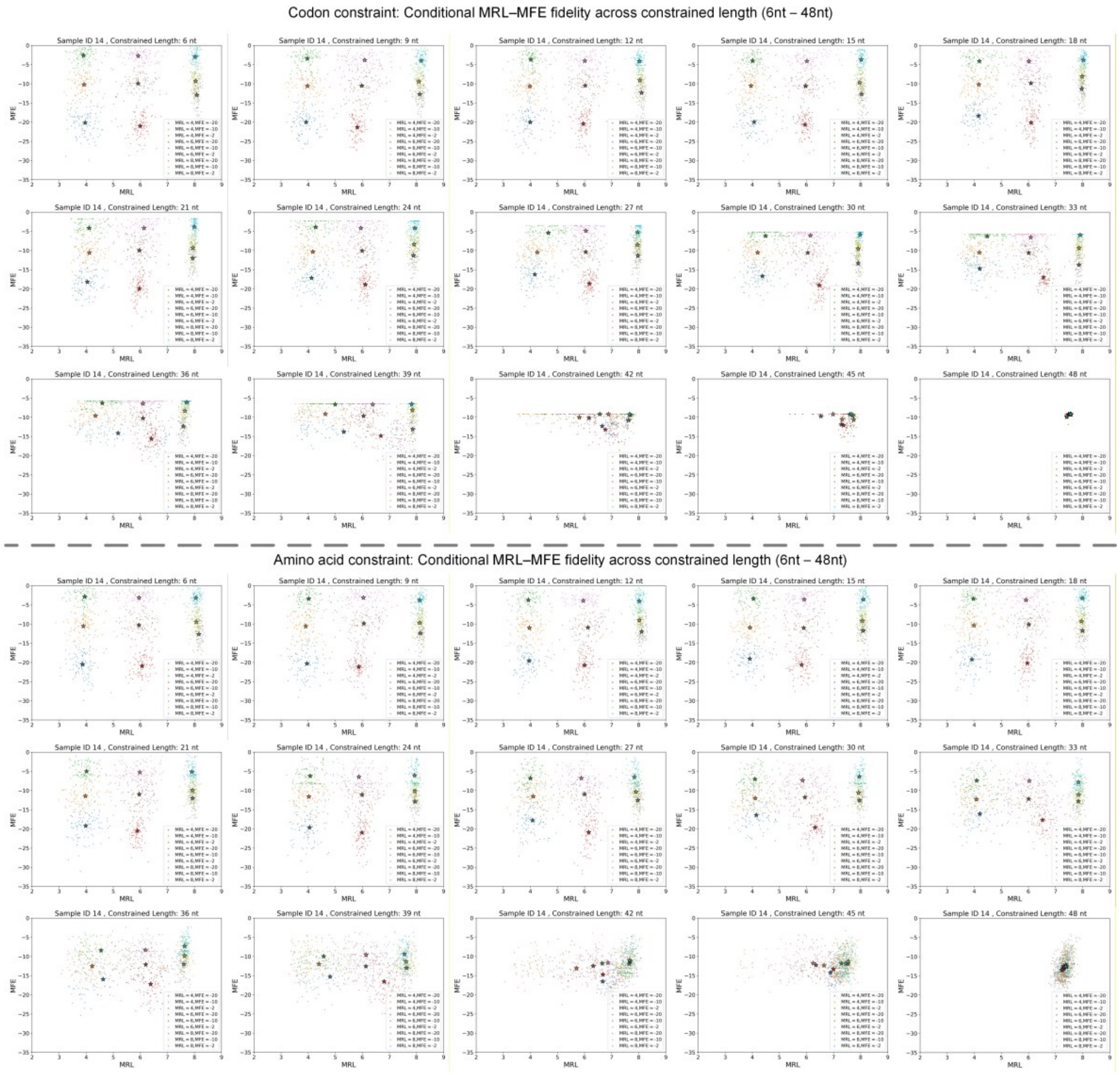

Figure S 9: MRL–MFE distribution of generated sequences under increasing constrained lengths under codon- and amino-acid-level constraints

Scatter plots show the distributions of generated sequences in the MRL–MFE plate for joint target values under codon-level constraints (top block) and amino-acid-level constraints (bottom block). Each panel corresponds to a fixed constrained CDS length, and colored point clouds represent different joint target pairs of MRL and MFE. Across all constraint lengths, generated sequences remain centered around their intended joint targets, indicating that the conditional fidelity is preserved even under strong local constraints.

As the constrained length increases, the distributions associated with each joint target progressively contract and ultimately converge toward compact clusters, reflecting reduced variability induced by stronger local constraints. While both constraint types exhibit similar convergence behavior, amino-acid-level constraints consistently maintain broader distributions around the convergence points than codon-level constraints, indicating higher residual sequence freedom under higher-level constraints.

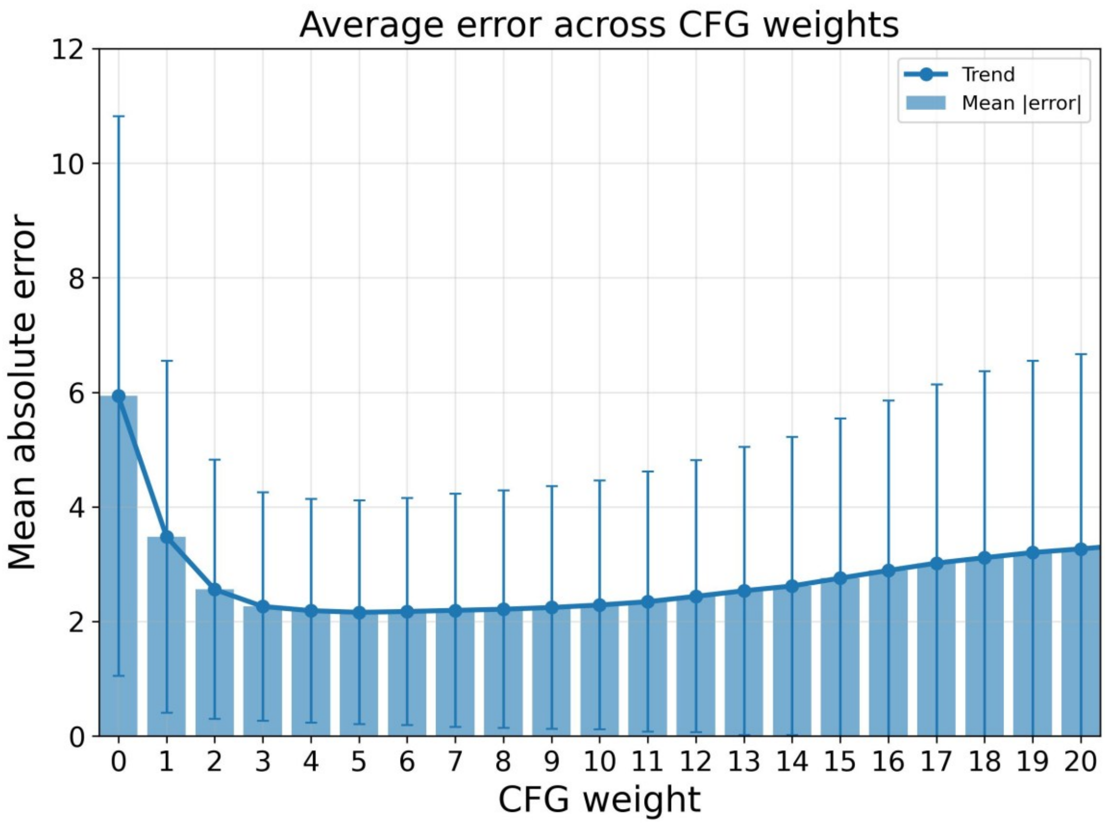

Figure S 10: Effect of classifier-free guidance weight on generation error

Mean absolute error averaged across generated samples is shown as a function of the classifier-free guidance (CFG) weight. Bars indicate the mean error at each CFG weight, and error bars represent the standard deviation across samples. The solid line highlights the overall trend.

Error decreases rapidly as the CFG weight increases from 0 to moderate values, reaching a minimum at intermediate weights, and gradually increases again at higher weights. This U-shaped relationship indicates a trade-off between insufficient conditioning at low CFG weights and over-guidance at high weights, suggesting that moderate CFG weights provide the best balance between conditional accuracy and sample variability.

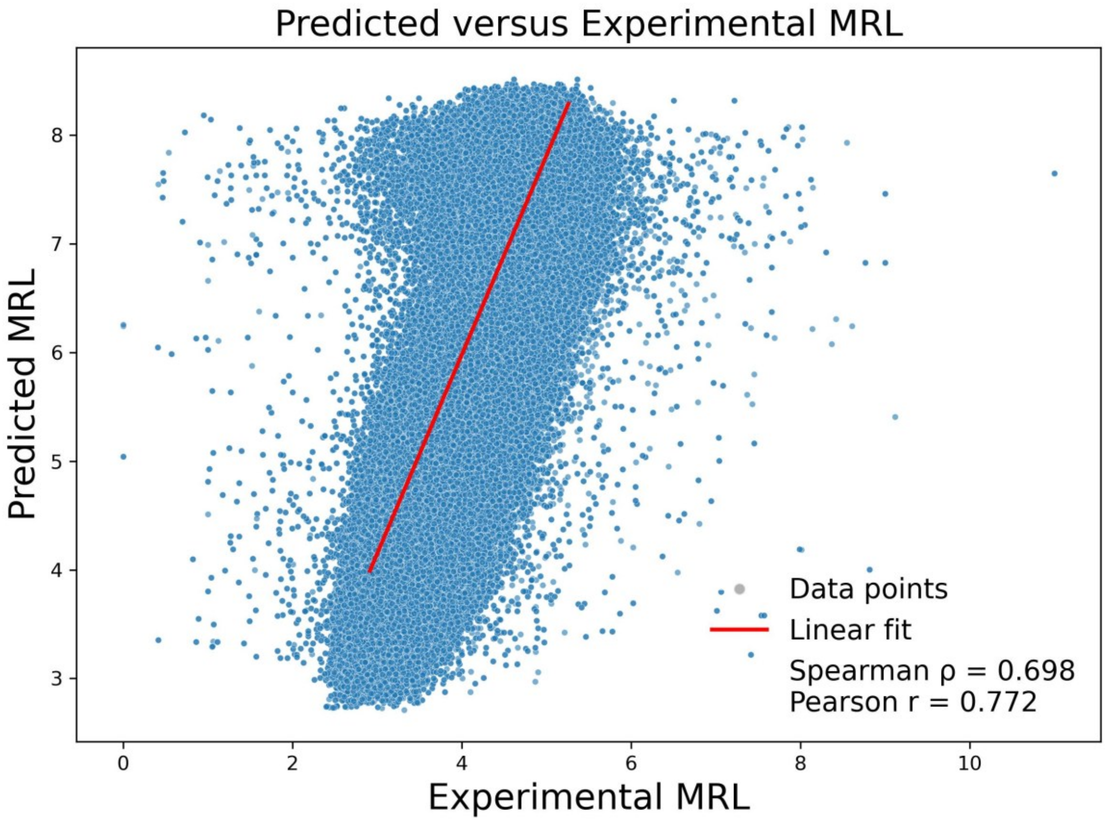

Figure S 11: Correlation between predicted and experimentally measured MRL

Scatter plot comparing UTR-LM–predicted MRL values used as training labels in this study with experimentally measured MRL obtained from polysome profiling of the same 5'UTR sequences. Each point corresponds to one sequence after length normalization to 50-nt. A linear regression line is shown in red. Predicted and experimental MRL exhibit a strong monotonic and linear correlation (Spearman ρ = 0.698, Pearson r = 0.772), indicating that predicted MRL captures relative expression trends in the experimental data, despite systematic deviations at the absolute scale.

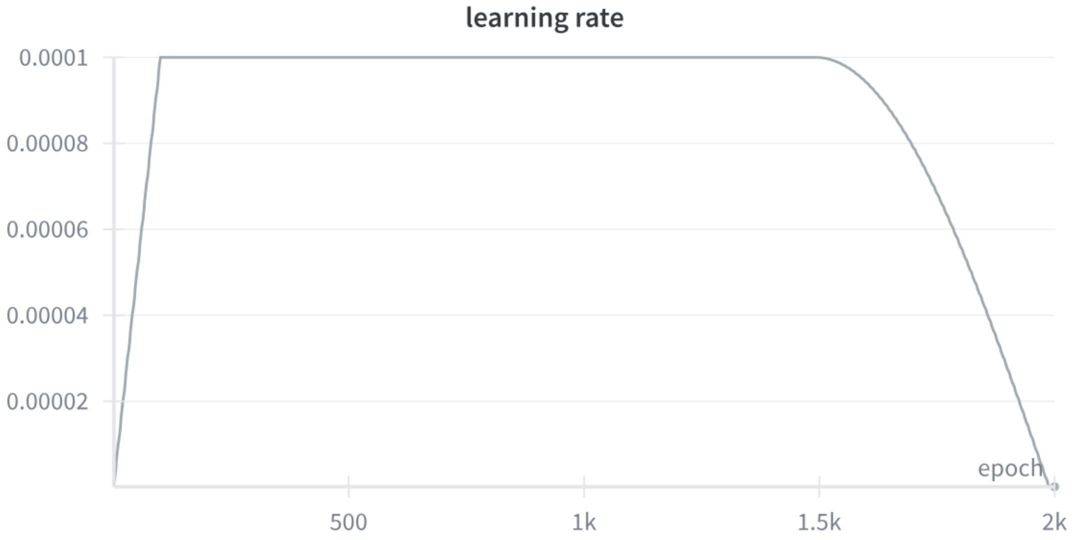

Figure S 12: Learning rate schedule used during training.

The learning rate follows a warmup–flatten–cosine decay schedule. After a brief warmup phase, the learning rate is held constant at its peak value for the majority of training before being gradually annealed in the final stage. The curve is shown as recorded during a representative training run.

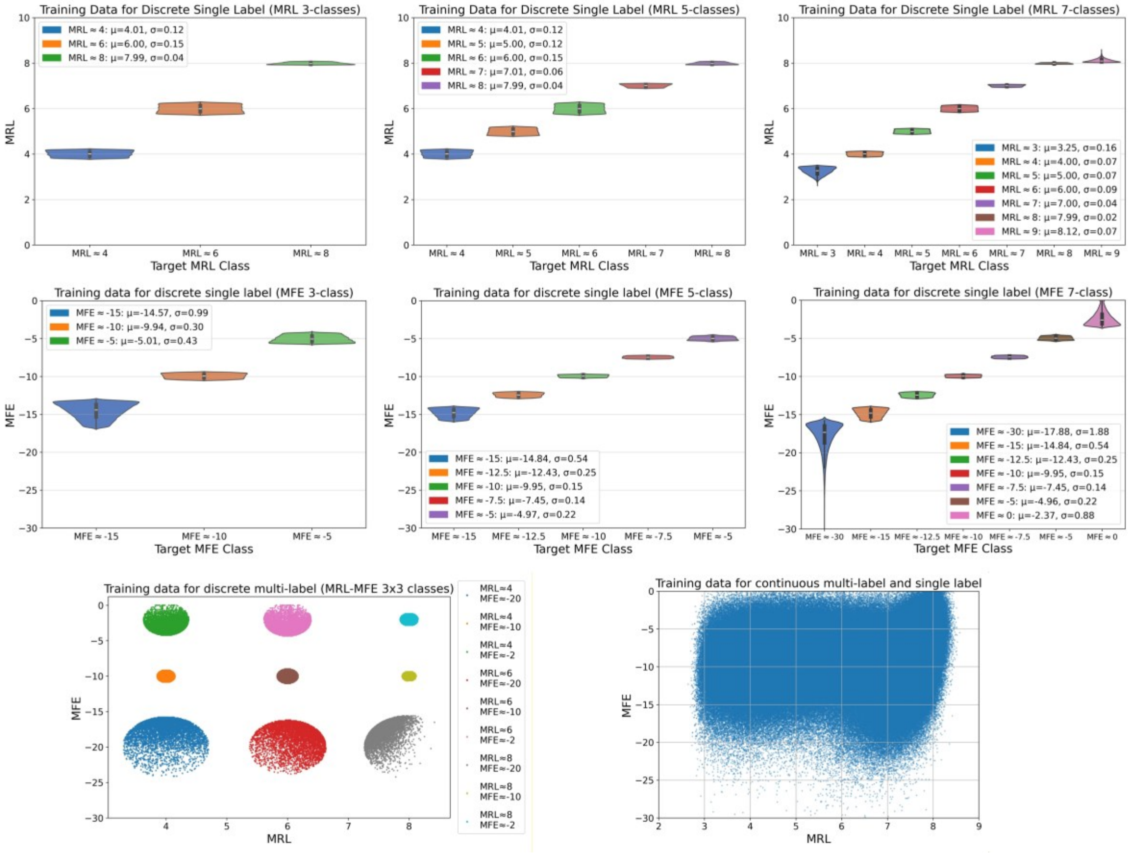

Figure S 13: Curation of training data for different conditional generation experiments

Training data distributions used for discrete single-label (MRL and MFE, 3/5/7 classes), discrete multi-label (MRL–MFE 3×3 classes), and continuous conditional generation settings. Violin and scatter plots illustrate the class-wise and joint distributions after data curation.

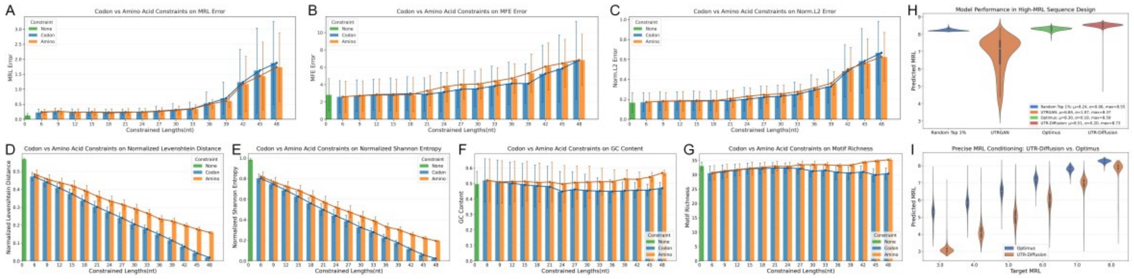

Figure S 14: Stress test under increasing contiguous CDS constraint length

To characterize sensitivity to progressively stronger local clamping, we evaluated five predefined CDS-like downstream reference segments and increased the contiguous constrained length from 6 to 48 nt under codon- and amino-acid-level constraints, while jointly conditioning on MRL and MFE. Because the analysis uses five predefined reference segments, it is treated as a targeted stress test rather than evidence of general robustness across CDS contexts.

Conditional error generally increased with constrained length, but the magnitude and onset of degradation depended on the joint target pair (Supplementary Figures S4 and S9). The codon- and amino-acid-level settings showed broadly similar length-dependent trends; the data did not support a uniform advantage of either constraint representation across all metrics and lengths.

Normalized Levenshtein distance and Shannon entropy decreased progressively as the constrained region lengthened (Supplementary Figures S5 and S6), whereas GC content remained comparatively stable and 3-mer motif richness showed only modest or target-dependent changes (Supplementary Figures S7 and S8). These results indicate a gradual trade-off between local sequence fixation, conditional fidelity, and generative diversity rather than an abrupt global collapse.

The experiment uses an artificial AUG within the 50-nt window and treats the downstream sequence as a simplified CDS-like proxy. It therefore probes model behavior under partial sequence clamping in synthetic UTR-like contexts and should not be interpreted as validation on endogenous 5′UTR–CDS junctions or coding regions.

### S2 Supplementary Tables

Table S 1 Summary of training data composition for different conditional generation experiments

| **Experiment Type** | **Label** | **Classes** | **Data per Class** | **Data per Class** |
|---|---|---|---|---|
| Discrete Single-label | MRL | 3-Class | 50,000 | 150,000 |
| | MRL | 5-Class | 50,000 | 250,000 |
| | MRL | 7-Class | 30,000 | 210,000 |
| | MFE | 3-Class | 100,000 | 300,000 |
| | MFE | 5-Class | 50,000 | 250,000 |
| | MFE | 7-Class | 25,000 | 175,000 |
| Discrete Double-label | MRL & MFE | 3×3 joint classes | 5,000 | 45,000 |
| Continuous Single-label | MRL | — | — | 967,647 |
| | MFE | — | — | 967,647 |
| Continuous Multi-label | MRL & MFE | — | — | 967,647 |

All local-constraint experiments, including the contiguous CDS-length stress test, were performed at sampling time using the pretrained continuous multi-label model, without additional fine-tuning.

Table S 2 Local coding constraint patterns and variants used in Figure 3

| Constraint class | Pattern family | Constrained nucleotide range(s) | Fixed or allowed identities | No. of variants |
|---|---|---|---|---|
| Codon/motif | Kozak | nt 20–29 | GCCACCAUGG; GCCGCCAUGG | 2 |
| | DRACH | DRACH: nt 12–16<br>Fixed AUG: nt 26–28 | DRACH variants (D = A/G/U, R = A/G, H = A/C/U):<br>AAACA, AAACC, AAACU, AGACA, AGACC, AGACU, GAACA, GAACC, GAACU, GGACA, GGACC, GGACU, UAACA, UAACC, UAACU, UGACA, UGACC, UGACU | 18 |
| | Kozak+DRACH | DRACH: nt 12–16<br>Kozak: nt 20–29 | All Cartesian combinations of the 18 DRACH sequences and the 2 Kozak sequences listed above | 36 |
| | Kozak+DRACH+triplet | Triplet: nt 6–8<br>DRACH: nt 12–16<br>Kozak: nt 20–29 | Triplets: UCA, AGU, CAG, GUA, ACU, UGA, CAU, GUC, AUC, CUA, GAC, UGC<br>Representative DRACH–Kozak pairs: UAACU–GCCACCAUGG (AU-heavy); GGACC–GCCGCCAUGG (GC-heavy); UGACC–GCCACCAUGG (balanced)<br>All 12 × 3 combinations | 36 |
| Amino acid | Single Amino | Amino acid: nt 38–40 | nt 38–40: D, A, H, G, V, N, K, R, L, W, F, E, Y, T, Q, I, C, P, S | 19 |
| | Double Amino | Amino acids: nt 35–37 and nt 41–43 | nt 35–37: D, A, H, V, S, N, K, R<br>nt 41–43: T, L, E, Y, G, Q, I, C<br>All 8 × 8 combinations | 64 |
| | Triple Amino | Amino acids: nt 32–34, nt 38–40, and nt 44–46 | nt 32–34: D, A, H, V<br>nt 38–40: S, N, K, R<br>nt 44–46: T, L, E, Y<br>All 4 × 4 × 4 combinations | 64 |
| | Quadra Amino | Amino acids: nt 29–31, nt 35–37, nt 41–43, and nt 47–49 | nt 29–31: D/A/H; nt 35–37: V/S/N; nt 41–43: K/R/T; nt 47–49: L/E/Y; all 3 × 3 × 3 × 3 combinations | 81 |

Note: Coordinates follow Figure 3; motifs use RNA notation and amino acids one-letter codes. Totals are Cartesian products; all amino-acid patterns fix AUG at nt 26–28.

**Table S 3 Reference peptide panel used for codon-adaptiveness-controlled generation in Figure 4**

| Reference ID | Gene | UniProt | Downstream peptide (7 aa) |
|---|---|---|---|
| Ref-01 | GAPDH | P04406 | GKVKVGV |
| Ref-02 | HPRT1 | P00492 | ATRSPGV |
| Ref-03 | HSPA8 | P11142 | SKGPAVG |
| Ref-04 | RPS9 | P46781 | PVARSWV |
| Ref-05 | RPL32 | P62910 | AALRPLV |
| Ref-06 | VCP | P55072 | ASGADSK |
| Ref-07 | PSMB4 | P28070 | EAFLGSR |
| Ref-08 | RPL13A | P40429 | AEVQVLV |
| Ref-09 | LMNA | P02545 | ETPSQRR |
| Ref-10 | VIM | P08670 | STRSVSS |

Note: All entries are from Homo sapiens (taxon 9606) and correspond to pep_21–pep_30 in Table S 4. The listed seven-residue peptide comprises the downstream amino acids constrained in Figure 4. The invariant initiation methionine encoded by the fixed AUG is omitted; the full constrained peptide is therefore M followed by the listed sequence (for example, Ref-01 corresponds to MGKVKVGV).

**Table S 4 Human reference peptide panel used for 5′ UTR–CDS junction benchmarking**

| Reference ID | Gene | UniProt | Downstream peptide (16 aa) | Benchmark coding target (10 aa) | log10 synonymous space (16 codons) |
|---|---|---|---|---|---|
| pep_01 | HSP90AA1 | P07900 | PEETQTQDQPMEEEEV | MPEETQTQDQ | 6.021 |
| pep_02 | NPM1 | P06748 | EDSMDMDMSPLRPQNY | MEDSMDMDMS | 6.424 |
| pep_03 | CALM1 | P0DP23 | ADQLTEEQIAEFKEAF | MADQLTEEQI | 6.674 |
| pep_04 | PSMA1 | P25786 | FRNQYDNDVTVWSPQG | MFRNQYDNDV | 6.975 |
| pep_05 | TPI1 | P60174 | APSRKFFVGGNWKMNG | MAPSRKFFVG | 6.975 |
| pep_06 | YWHAZ | P63104 | DKNELVQKAKLAEQAE | MDKNELVQKA | 6.975 |
| pep_07 | EEF1A1 | P68104 | GKEKTHINIVVIGHVD | MGKEKTHINI | 7.151 |
| pep_08 | GPI | P06744 | AALTRDPQFQKLQQWY | MAALTRDPQF | 7.151 |
| pep_09 | TUBB | P07437 | REIVHIQAGQCGNQIG | MREIVHIQAG | 7.327 |
| pep_10 | ACTB | P60709 | DDDIAALVVDNGSGMC | MDDDIAALVV | 7.452 |
| pep_11 | LDHA | P00338 | ATLKDQLIYNLLKEEQ | MATLKDQLIY | 7.503 |
| pep_12 | ALDOA | P04075 | PYQYPALTPEQKKELS | MPYQYPALTP | 7.753 |
| pep_13 | PPIA | P62937 | VNPTVFFDIAVDGEPL | MVNPTVFFDI | 7.878 |
| pep_14 | PSMB2 | P49721 | EYLIGIQGPDYLVASD | MEYLIGIQGP | 8.105 |
| pep_15 | TUBA1B | P68363 | RECISIHVGQAGVQIG | MRECISIHVG | 8.105 |
| pep_16 | HMBS | P08397 | SGNGNAAATAEENSPK | MSGNGNAAAT | 8.179 |
| pep_17 | CFL1 | P23528 | ASGVAVSDGVIKVFND | MASGVAVSDG | 8.355 |
| pep_18 | RPS18 | P62269 | SLVIPEKFQHILRVLN | MSLVIPEKFQ | 8.457 |
| pep_19 | PCNA | P12004 | FEARLVQGSILKKVLE | MFEARLVQGS | 8.582 |
| pep_20 | PGK1 | P00558 | SLSNKLTLDKLDVKGK | MSLSNKLTLD | 8.582 |
| pep_21 | GAPDH | P04406 | GKVKVGVNGFGRIGRL | MGKVKVGVNG | 8.832 |
| pep_22 | HPRT1 | P00492 | ATRSPGVVISDDEPGY | MATRSPGVVI | 8.832 |

| Reference ID | Gene | UniProt | Downstream peptide (16 aa) | Benchmark coding target (10 aa) | log10 synonymous space (16 codons) |
|---|---|---|---|---|---|
| pep_23 | HSPA8 | P11142 | SKGPAVGIDLGTTYSC | MSKGPAVGID | 8.832 |
| pep_24 | RPS9 | P46781 | PVARSWVCRKTYVTPR | MPVARSWVCR | 8.832 |
| pep_25 | RPL32 | P62910 | AALRPLVKPKIVKKRT | MAALRPLVKP | 9.008 |
| pep_26 | VCP | P55072 | ASGADSKGDDLSTAIL | MASGADSKGD | 9.184 |
| pep_27 | PSMB4 | P28070 | EAFLGSRSGLWAGGPA | MEAFLGSRSG | 9.309 |
| pep_28 | RPL13A | P40429 | AEVQVLVLDGRGHLLG | MAEVQVLVLD | 9.309 |
| pep_29 | LMNA | P02545 | ETPSQRRATRSGAQAS | METPSQRRAT | 9.786 |
| pep_30 | VIM | P08670 | STRSVSSSSYRRMFGG | MSTRSVSSSS | 10.014 |

Note: All entries are from Homo sapiens (taxon 9606). The 16-aa downstream peptide is listed as obtained from the source protein. Benchmark 2 uses the fixed AUG-encoded methionine followed by the first nine downstream residues, yielding the 10-aa coding target shown. log10 synonymous space denotes log10 of the synonymous coding-space size for the full 16-aa downstream peptide.

S3 **Supplementary Methods**

### S3.1 Evaluation Measures and Normalization

Evaluation measures were selected according to the objective of each analysis. All target-dependent measures were calculated separately for each MRL–MFE target class. For a class c containing N generated sequences, $q_c$ denotes the requested value of indicator q ∈ {MRL, MFE}, $\hat{q}_{c,i}$ denotes the corresponding predicted value for sequence i, and $\bar{q}_c$ denotes the class mean. Target-wise distributions were visualized directly, whereas scalar summaries were calculated as described below.

**Conditional fidelity.** For each indicator, we summarized the class mean and its signed deviation from the requested target. Where reported, mean absolute error (MAE) was calculated across generated sequences:

$$\bar{q}_c = (1/N)\, \Sigma_i\, \hat{q}_{c,i}, \qquad \Delta q_c = \bar{q}_c - q_c, \qquad MAE_q(c) = (1/N)\, \Sigma_i\, |\hat{q}_{c,i} - q_c|.$$

For joint MRL–MFE fidelity, the signed class-mean errors were combined after scaling each axis by its evaluated target range:

$$E_{joint}(c) = \sqrt{[(\Delta MRL_c/R_{MRL})^2 + (\Delta MFE_c/R_{MFE})^2]}.$$

Here, $R_{MRL} = \max_c(MRL_c) - \min_c(MRL_c)$ and $R_{MFE} = \max_c(MFE_c) - \min_c(MFE_c)$. When an explicit evaluation range was supplied, that range was used instead. In the vector-field plots, arrow direction represents the unnormalized signed errors ($\Delta MRL_c$, $\Delta MFE_c$), whereas arrow length and heatmap color represent $E_{joint}(c)$.

**Generative diversity.** Position-wise nucleotide uncertainty was quantified by Shannon entropy. For nucleotide probabilities $p_{l,b}$ at position l and b ∈ {A, C, G, U}, the entropy at that position and its sequence-level normalized mean were:

$$H_l = -\Sigma_b\, p_{l,b} \log_2(p_{l,b}), \qquad H_{norm} = [1/(L \log_2 4)]\, \Sigma_l\, H_l.$$

At amino-acid-constrained sites, codon-level entropy was calculated across the permitted synonymous codons and normalized by $\log_2|Syn(a)|$, where Syn(a) is the synonymous-codon set for the constrained amino acid a. Sites with only one permitted codon were assigned a normalized codon entropy of 0.

Thus, $H_{norm}$ ranges from 0 for complete positional conservation to 1 for an equal nucleotide distribution at every position. Whole-sequence diversity was quantified using the mean pairwise Levenshtein distance normalized by sequence length:

$$D_{Lev} = [2/(N(N-1)L)]\, \Sigma_{i<j}\, d_{Lev}(s_i, s_j),$$

where d_Lev is the minimum number of single-nucleotide insertions, deletions, and substitutions required to transform one sequence into another, and L is the sequence length. All unique unordered sequence pairs were included, and self-comparisons were excluded.

**Sequence composition and coding adaptation.** For each sequence s_i, GC content was calculated as

$$GC(s\_i) = [n\_G(s\_i) + n\_C(s\_i)]/L.$$

The 3-mer motif richness of a sequence was defined as the number of distinct valid RNA 3-mers observed in that sequence:

$$R_3(s\_i) = |\{s\_i[l{:}l{+}3] : 1 \le l \le L{-}2\}|.$$

GC content and 3-mer richness were summarized by their mean and standard deviation across sequences in each target class. Natural training sequences and length-matched random sequences were used as reference distributions; random sequences were generated by independently sampling A, C, G, and U with equal probability at every position.

For experiments with a specified peptide, codon adaptation index (CAI) was calculated from the relative adaptiveness w(c_j) of each evaluated synonymous codon c_j:

$$w(c\_j) = f(c\_j|a\_j)/\max\_\{c\in Syn(a\_j)\}\ f(c|a\_j), \qquad CAI = \exp[(1/K)\ \Sigma_j \log w(c\_j)].$$

Here, f(c|a) is the human codon-usage frequency of codon c for amino acid a, Syn(a) is the synonymous-codon set for a, and K is the number of evaluated codons after experiment-specific exclusions. The fixed AUG initiation codon was excluded from CAI calculation. CAI7 was calculated over the seven downstream codons in Figure 4, whereas CAI9 was calculated over the nine downstream codons in Figure 5. Unless otherwise specified, plotted curves and heatmaps report target-class means; error bars show one standard deviation when present.

**Aggregation in Figure 4.** In Figure 4A, each point represents the mean predicted MRL–MFE pair across 100 generated sequences for one reference peptide and one codon-adaptiveness setting within a target class; each star is the mean of the 30 peptide-by-adaptiveness point estimates in that class. In Figure 4B, bars and error bars show the mean ± one standard deviation across the nine MRL–MFE target-condition means, and the overlaid points show the corresponding condition-specific means.

**Aggregation in Figure 5.** In Figure 5C, each small point represents the mean of 100 generated sequences for one reference peptide. The large marker and error bar show the mean ± one standard deviation across the 30 reference-peptide means for each design method. Thus, generated sequences within a reference-peptide batch were summarized before comparison across reference peptides.

#### S3.2 Diffusion Framework

We adopted the Denoised Diffusion Probabilistic Model (DDPM) [Ho et al., 2020] framework to generate RNA sequences. The model consists of two complementary processes: a forward diffusion process that gradually corrupts a clean sequence by adding Gaussian noise, and a reverse denoising process that iteratively removes noise to sample from the data distribution.

The diffusion process gradually perturbs the one-hot representation $x_t \in \mathbb{R}$^{4×L} of clean sequence $x_0$ by adding Gaussian noise at each step, so that after T steps the representation x_T becomes nearly Gaussian noise. Each diffusion step is defined as:

$$q(x_t \mid x_{t-1}) \sim \mathcal{N}(\sqrt{(1-\beta_t)}\, x_{t-1}, \beta_t I) \quad \text{(S1)}$$

where $\alpha_t = 1 - \beta_t$ and $\bar{\alpha}_t = \prod_{s=1}^{t} \alpha_s$. Using these definitions, $x_t$ admits a closed-form expression:

$$x_t = \sqrt{\bar{\alpha}_t}\, x_0 + \sqrt{(1-\bar{\alpha}_t)}\, \varepsilon, \qquad \varepsilon \sim \mathcal{N}(0, I) \quad \text{(S2)}$$

The reverse process starts from Gaussian noise x_T and, at each step t, predicts the noise component to be removed from $x_t$, gradually inferring the underlying sample $x_0$. Each reverse step is defined as:

$$p\theta(x_{t-1} \mid x_t) \sim \mathcal{N}(\mu\theta(x_t, t), \Sigma\theta(x_t, t)) \quad \text{(S3)}$$

Here, the variance term $\Sigma\theta(x_t, t)$ can be reparameterized using time-dependent constants:

$$\Sigma\theta(x_t, t) = \sigma_t^2 = [(1-\bar{\alpha}_{t-1})/(1-\bar{\alpha}_t)]\, \beta_t \quad \text{(S4)}$$

The mean term $\mu\theta(x_t, t)$ can be expressed in terms of the predicted noise $\varepsilon\theta(x_t, t)$:

$$\mu\theta(x_t, t) = (1/\sqrt{\alpha_t})\, [x_t - \beta_t/\sqrt{(1-\bar{\alpha}_t)}\, \varepsilon\theta(x_t, t)] \quad \text{(S5)}$$

This parameterization emphasizes that the neural network is trained to predict the added noise $\varepsilon\theta(x_t, t)$, which enables recovery of the original sequence $x_0$. Accordingly, the training objective minimizes the discrepancy between the true noise ε and the predicted noise:

$$\mathcal{L}(\theta) = E_{t, x0, \varepsilon}\, [\|\varepsilon - \varepsilon\theta(\sqrt{\bar{\alpha}_t}\, x_0 + \sqrt{(1-\bar{\alpha}_t)}\, \varepsilon, t)\|^2] \quad (6)$$

This noise prediction objective corresponds to the simplified loss proposed by DDPM, which is widely adopted in diffusion-based generative models.

#### S3.3 Data Curation and Dataset Splitting

We obtained a total of 967,647 unique 50-nt 5′ UTR sequences after preprocessing, as described in Section 2.1. While this full dataset was used for several experiments, not all experiments relied on the complete collection. In particular, for a subset of experiments, an additional data curation step was performed prior to model training.

This curation step was applied to discrete-label generation experiments, including both discrete single-label and discrete multi-label settings, in order to construct well-defined class-specific training subsets. Because the original MRL and MFE labels are continuously distributed, directly assigning sequences to discrete classes using fixed bin boundaries can introduce substantial ambiguity near class edges. To address this issue, we adopted a label-distance–based selection strategy. For each target discrete label value, sequences were ranked according to their absolute distance to the target value in the label space, and the top-ranked sequences were selected to form the corresponding class-specific subset. Depending on the experiment, the number of selected sequences per class (N) was set to 25,000, 50,000, or 100,000.

Importantly, this curation procedure was based solely on label values and was performed prior to any model training, without reference to sequence features or model outputs. The resulting curated subsets were used exclusively for discrete-label generation experiments. In contrast, continuous-label generation experiments were conducted using the full dataset without additional filtering. The datasets used for each experiment are summarized in Table S 1. The label distributions of the curated training datasets used for all discrete-label generation experiments are shown in Figure S 12.

For all experiments, the selected datasets were randomly split into training and validation sets using a fixed ratio of 95% and 5%, respectively.

#### S3.4 Training Configuration

All experiments were trained using a unified training configuration, regardless of whether the task involved discrete-label or continuous-label generation, or whether codon/amino-acid constraints were enforced.

Model parameters were optimized using the AdamW optimizer with a peak learning rate of $1 \times 10^{-4}$, weight decay of $2 \times 10^{-4}$, and momentum parameters $\beta_1 = 0.9$ and $\beta_2 =$

0.95. All models were trained for 2000 epochs with a batch size of 6000. Model performance was monitored on a held-out validation set every five epochs.

A structured learning rate schedule was applied throughout training. The total number of training steps was defined as the product of the number of training batches per epoch and the total number of training epochs. The learning rate was linearly increased from zero during an initial warmup phase covering 5% of the total training steps. This was followed by a flattened phase, during which the learning rate was held constant at its peak value for 70% of the training steps. In the remaining phase, the learning rate was gradually decayed using a cosine schedule, with a lower bound to prevent vanishing learning rates. A representative example of the learning rate schedule recorded during training is shown in Figure S 12.

To stabilize training and improve sampling quality, an exponential moving average (EMA) of model parameters was maintained with a decay rate of 0.995. The EMA model was used for validation and sequence generation unless otherwise specified.

Discrete-label experiments were trained on a single NVIDIA H100 GPU, whereas continuous-label experiments were trained using distributed data-parallel execution across four NVIDIA H100 GPUs to accommodate the larger training dataset and reduce wall-clock training time. No changes were made to the model architecture or training hyperparameters between these settings.

#### S3.5 UNet Architecture

The denoising network is implemented as a compact U-Net architecture with three resolution levels. Two down-sampling operations are applied along the sequence width dimension, progressively reducing the feature width from 4 to 1, followed by a symmetric up-sampling path with skip connections between corresponding levels.

The model operates on a single-channel input tensor of shape (1, 64, 4). To accommodate the fixed network architecture, the original 50-nt sequence is padded with eight positions on both the 5′ and 3′ sides prior to input, resulting in an effective sequence length of 64. This padding is applied uniformly across all experiments and does not introduce additional sequence information.

Temporal information is incorporated through a diffusion timestep embedding, which is injected into residual blocks across multiple resolution levels. For conditional generation tasks, conditioning labels are projected into the temporal embedding space and added element-wise to the diffusion time embedding. This fused temporal representation is used to condition residual blocks across multiple spatial resolutions in the U-Net, ensuring consistent injection of both temporal and conditional information throughout the network.

Unless otherwise specified, the same U-Net architecture is used for all experiments.

#### S3.6 Diffusion Model Parameters

All experiments are based on a denoising diffusion probabilistic model (DDPM) formulation. Given an input sequence representation $x_0$, the forward diffusion process progressively adds Gaussian noise over T = 200 timesteps according to a fixed variance schedule. Specifically, a linear $\beta_t$ schedule is used, with $\beta_t$ increasing linearly from a small initial value to a final value of β_T = 0.01. The corresponding $\alpha_t$ and cumulative-product $\bar{\alpha}_t$ terms are precomputed and used for both training and sampling.

During training, the model is optimized to predict the added noise ε at a randomly sampled timestep t, following the standard DDPM objective. No modifications are made to the diffusion objective between discrete-label and continuous-label settings. The same noise schedule, timestep count, and loss formulation are used across all experiments.

The distinction between discrete-label and continuous-label experiments lies solely in the form of the conditioning variables provided to the model. Discrete-label experiments use categorical class labels, whereas continuous-label experiments condition on continuous-valued or multi-dimensional label vectors. Importantly, these differences do not alter the underlying diffusion process or noise parameterization.

#### S3.7 Classifier-free Guidance Strategy

Classifier-free guidance (CFG) is employed during sampling to control the strength of conditional generation by interpolating between conditional and unconditional denoising predictions using a guidance weight. During training, conditional information is randomly masked with a fixed probability (unconditional rate = 0.2), enabling a single model to learn both conditional and unconditional denoising behaviors without relying on an external classifier.

Importantly, the “unconditional” representation here is not truly unconditional, but rather a mixture of all classes. As a result, CFG effectively pushes a specific class away from the global class mixture (even though the mixture also contains itself), thereby reinforcing its distinction from all other classes.

To examine the effect of the CFG weight, we performed a sweep over a range of guidance weights and evaluated the resulting generation error. As shown in Figure S 10, the absolute mean error exhibits a characteristic U-shaped relationship with respect to the

CFG weight. Error decreases rapidly as the CFG weight increases from zero to moderate values, reaches a minimum at intermediate weights, and gradually increases again at higher weights. This trend reflects the trade-off between insufficient conditioning at low guidance strengths and over-guidance at high values, which can degrade sample quality and increase variability.

In all experiments reported in this study, a fixed CFG weight of 1.0 was used. This choice was made to ensure consistency and comparability across different experimental settings and conditioning tasks. Importantly, for the primary conditional targets considered in this work (e.g., MRL), a guidance weight of 1 already yields strong conditional accuracy, as indicated by the low generation error observed in the sweep analysis.

We note that the optimal CFG weight is generally task- and label-dependent. Prior work has reported that a guidance weight of approximately 4 is optimal for text-conditioned image generation tasks. In our experiments, the optimal guidance weight for RNA sequence generation conditioned on MFE is observed to be around 5, as shown in Figure S 10. Therefore, while a fixed guidance weight was adopted here for methodological consistency, further task-specific tuning of both the guidance weight and the unconditional rate represents a straightforward avenue for improving conditional fidelity in future work.

Timestep embedding: The scalar diffusion timestep t is first encoded using a sinusoidal positional representation, followed by a two-layer MLP with GELU activation. This produces a time-dependent feature vector that is injected into each UNet block.

Discrete-label embedding: For categorical labels (e.g., 3-class MRL label), conditioning is implemented through a standard embedding lookup table that maps each label to a 200-dimensional vector.

Continuous-label embedding: For quantitative indicators such as MRL or MFE metrics, the label values are projected through a two- layer MLP with SiLU activation to obtain a 200-dimensional vector.

These embeddings are added to the intermediate feature maps within the UNet, enabling the model to incorporate both timestep and expression information at each diffusion and reverse step.
